# Temozolomide-associated inflammatory-repair remodeling in recurrent glioblastoma exposes a ROCK-linked therapeutic vulnerability

**DOI:** 10.64898/2026.09.20.752931

**Authors:** Rafal Chojak, Umme H. Faisal, Priya U. Kumthekar, Noah B. Drewes, Lara Koutah, Jessica Jia, Jillyn R. Turunen, Maeve C. O’Shea, Victoria Falkowski, Ayaan Akhtar, Shutong Wang, Justin Chen, Shanti Ahmed, Hasaan Kazi, Brenda Auffinger, Pouya Jamshidi, Katy McCortney, Alicia Marie Catezone, Matthew C. Tate, Roger Stupp, Maciej S. Lesniak, Jason Miska, Karan S. Dixit, Ditte Primdahl, Dieter H. Heiland, Adam M. Sonabend, Atique U. Ahmed

## Abstract

**Background:** Glioblastoma (GBM) adapts to therapy through coordinated malignant-cell and microenvironmental responses, but the mechanisms linking treatment-associated inflammation to tumor-cell phenotypic plasticity remain poorly defined. Here, we investigated whether preoperative temozolomide exposure is associated with inflammatory-repair remodeling and a ROCK-linked therapeutic vulnerability in recurrent GBM using a window-of-opportunity clinical trial cohort.

**Methods:** We integrated patient-resolved single-cell transcriptomics, spatial RNA profiling and multiplex protein imaging with experimental perturbations and orthotopic glioblastoma models. In the presurgical window-of-opportunity subgroup of NCT05236036, four patients with recurrent glioblastoma received additional preoperative temozolomide (TMZ; 150 mg/m2/day for 5 days) before resection and were compared with three recurrent comparators who did not receive additional preoperative TMZ.

**Results:** Across seven tumors, malignant-cell inflammatory activity covaried with integrin-binding and wound-healing programs (median partial ρ = 0.45 and 0.42, respectively). Tumors exposed to preoperative TMZ showed enrichment of inflammatory, chemokine, adhesion and wound-healing programs (GSEA q < 0.05), with the direction of enrichment preserved in all leave-one-patient-out analyses. In two tumors exposed to preoperative TMZ, CXCL12 localized to vascular territories, whereas chemokine and wound-repair programs increased near injury-reference regions. Across seven specimens analyzed by multiplex protein imaging, MYL9 abundance correlated with local CXCL12 (ρ = 0.61), phosphorylated MYPT1 (ρ = 0.48) and nuclear phosphorylated STAT3 (ρ = 0.45), with positive associations in every specimen. Experimentally, TMZ increased CXCL8 and reactive-state markers and enhanced subsequent scratch closure, while CXCL8 and CXCL12 increased MLC2 phosphorylation. Fasudil attenuated TNF-NF-κB, inflammatory-response and IFN-γ-response programs in TMZ-treated cells and reduced scratch closure and p-MLC2 in complementary assays. Across three orthotopic models, fasudil plus TMZ prolonged survival versus TMZ alone (model-stratified HR, 0.21; 95% CI, 0.07-0.61; P = 0.001).

**Conclusions:** These findings link treatment-associated inflammatory-repair programs with cytoskeletal remodeling and support further translational evaluation of fasudil plus TMZ in glioblastoma.

## INTRODUCTION

Temozolomide (TMZ), an alkylating agent used as part of standard-of-care therapy for glioblastoma, improves survival when combined with radiotherapy, but disease progression remains nearly universal [1]. The behavior of cells that survive treatment is therefore central to understanding therapeutic failure. Cytotoxic exposure may alter cellular programs as well as reduce tumor burden, potentially changing how residual cells interact with the surrounding brain. Identifying which treatment-associated responses support persistence, and which can be pharmacologically interrupted, could inform combinations that extend the benefit of standard therapy.

Glioblastoma cells occupy plastic transcriptional states shaped by genetic and microenvironmental influences [2]. Inflammatory signaling is one such influence. TNF–NF-κB signaling promotes mesenchymal differentiation and radioresistance, whereas macrophage-derived oncostatin M can induce mesenchymal-like states through OSMR/LIFR–GP130 signaling [3,4]. Longitudinal studies of glioblastoma across diagnosis and recurrence have documented substantial genetic, transcriptional and microenvironmental evolution [5,6]. Recurrent tumors therefore reflect the combined effects of prior therapy, clonal selection and changes in the tumor microenvironment, complicating attribution of molecular features to a specific treatment. Analysis of tissue obtained during defined treatment exposure provides an opportunity to identify programs associated with recent therapy and determine how they are spatially organized within the tumor.

Spatial studies have placed reactive glioblastoma programs within structured vascular, immune and hypoxic environments [7,8]. Perivascular CXCL8 (IL-8) signaling can regulate glioblastoma stem-cell phenotypes [9], while injury-response programs and tumor-associated axonal damage connect the tumor ecosystem to tissue-repair biology [10,11]. Experimental studies have also linked TMZ exposure to chemokine remodeling and enhanced invasiveness, including CXCL8/CXCR2-associated cellular plasticity [12–14]. Together, these findings suggest that treatment-associated inflammatory and tissue-repair programs may converge on cytoskeletal remodeling, but how these programs are coupled in human glioblastoma during recent TMZ exposure remains poorly defined.

ROCK kinases regulate actomyosin organization and provide a candidate point of intervention. Fasudil has shown anti-invasive activity in experimental glioblastoma, and enhanced TMZ sensitivity has been reported in resistant glioma models, providing a rationale for evaluating ROCK inhibition in this setting [15,16].

We therefore asked whether inflammatory–repair programs associated with preoperative TMZ exposure are linked to therapeutically targetable cytoskeletal remodeling in recurrent glioblastoma. Presurgical TMZ was associated with inflammatory, chemokine, adhesion and wound-healing programs, while spatial and multiplex analyses linked vascular CXCL12 and repair-associated features to cytoskeletal signaling. In complementary experimental models, TMZ induced inflammatory and reactive phenotypes, CXCL8 and CXCL12 increased MLC2 phosphorylation, and the ROCK inhibitor fasudil suppressed inflammatory and cytoskeletal responses. Together, these findings nominate ROCK-associated actomyosin signaling as a potential therapeutic vulnerability in TMZ-exposed recurrent glioblastoma, which we further evaluated in orthotopic models.

## RESULTS

### Temozolomide exposure is associated with inflammatory and tissue-repair programs in recurrent human glioblastoma

We integrated malignant-cell transcriptomes, spatial RNA profiling and multiplex protein imaging to examine inflammatory, tissue-repair and cytoskeletal programs in recurrent glioblastoma. The single-cell cohort included four patients who received additional preoperative TMZ and three recurrent comparators without additional preoperative TMZ; all patients had previously received TMZ and radiotherapy (**Fig. 1A**). The annotated single-cell dataset comprised 165,625 cells: 45,915 from recurrent comparators and 119,710 from patients receiving additional preoperative TMZ. Broad-lineage and cell-type annotations are summarized in **Supplementary Fig. 1**, with corresponding marker-expression profiles shown in **Supplementary Fig. 2**. Analyses included 32,991 malignant cells selected using a joint RNA-inferred chromosome 7 gain/chromosome 10 loss criterion (**Fig. 1B; Supplementary Fig. 1A,D**).

**Figure 1.**
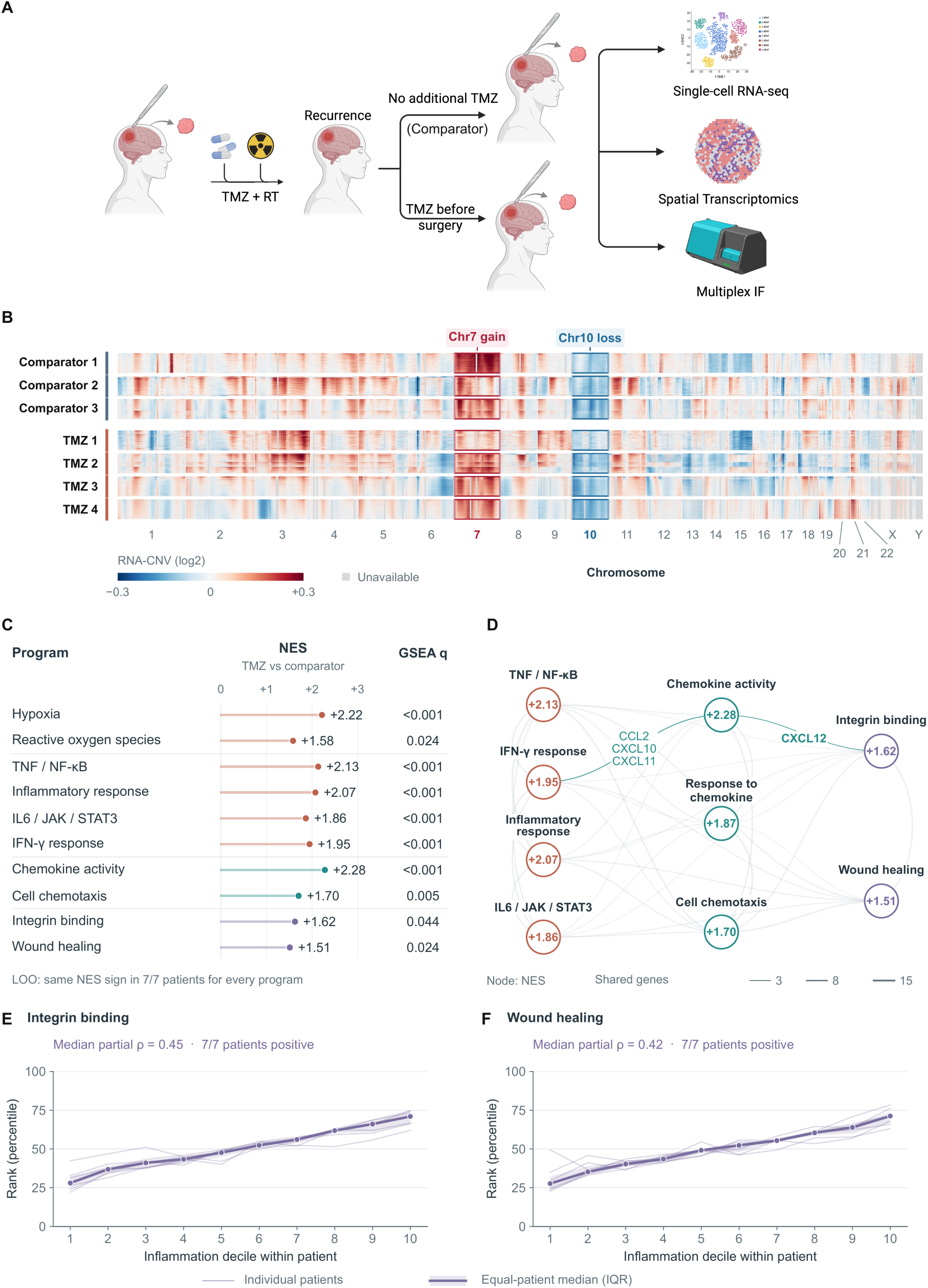
Coordinated inflammatory–repair transcription in recurrent glioblastoma. **A**, Study design: three recurrent comparators without additional preoperative TMZ and four patients receiving additional preoperative TMZ; all previously received TMZ and radiotherapy. **B**, RNA-inferred copy-number profiles for 32,991 cells selected using joint chromosome 7 gain/chromosome 10 loss. Columns represent 535 genomic bins and equal-height bands represent individual patients; colors saturate at ±0.3 and gray denotes unavailable values. **C**, Selected NES from patient-pseudobulk limma–voom-ranked GSEA. q values retain Benjamini–Hochberg correction within the corresponding full Hallmark or pooled Gene Ontology testing family. All displayed NES directions were preserved in seven leave-one-patient-out analyses; this denotes directional stability, not significance in every omission analysis. **D**, Leading-edge overlap network: node values indicate NES and edge widths indicate shared-gene counts, not directed signaling. **E**,**F**, Integrin-binding and wound-healing ranks across within-patient inflammation deciles, using disjoint signatures and technical-covariate adjustment. Thin lines represent patients; thick lines and shading indicate equal-patient medians and interquartile ranges. Annotations summarize patient-specific partial Spearman correlations. Deciles were defined separately for each signature pair.

Gene-set enrichment analysis (GSEA) of genes ranked by patient-level pseudobulk differential expression identified enrichment of hypoxia, TNF–NF-κB, inflammatory-response and chemokine-activity programs in the additional-TMZ group (normalized enrichment scores [NES], 2.22, 2.13, 2.07 and 2.28, respectively; q < 0.001 each). Cell chemotaxis, integrin binding and wound healing were also enriched (NES, 1.70, 1.62 and 1.51; q = 0.005, 0.044 and 0.024; **Fig. 1C**). Leading-edge overlap connected inflammatory, chemokine and repair annotations (**Fig. 1D**). All displayed enrichment directions were preserved across seven leave-one-patient-out analyses.

Within individual tumors, inflammation correlated positively with integrin-binding and wound-healing programs using disjoint signatures and technical-covariate adjustment (median partial Spearman ρ = 0.45 and 0.42; positive in all seven patients; **Fig. 1E,F**). These within-patient associations complement the between-group comparison and support coordinated inflammatory–repair transcription. Among the shared leading-edge genes, CXCL12 linked the chemokine-activity and integrin-binding programs (**Fig. 1D**), motivating us to examine its spatial organization in human tissue.

### Vascular CXCL12 and injury-associated chemokine–repair programs define distinct spatial niches in TMZ-exposed glioblastoma

Spatial RNA profiling of two TMZ-exposed IDH-wildtype glioblastoma sections, 502A2 and 504A2, localized chemokine and repair features to distinct tissue contexts (**Fig. 2A**). Exploratory block-level associations included MYL9–STAT3 in 502A2 (ρ = 0.30; conditional 95% spatial-bootstrap interval, 0.23–0.36) and CD44–VIM in 504A2 (ρ = 0.40; 0.33–0.47; **Fig. 2B**).

**Figure 2.**
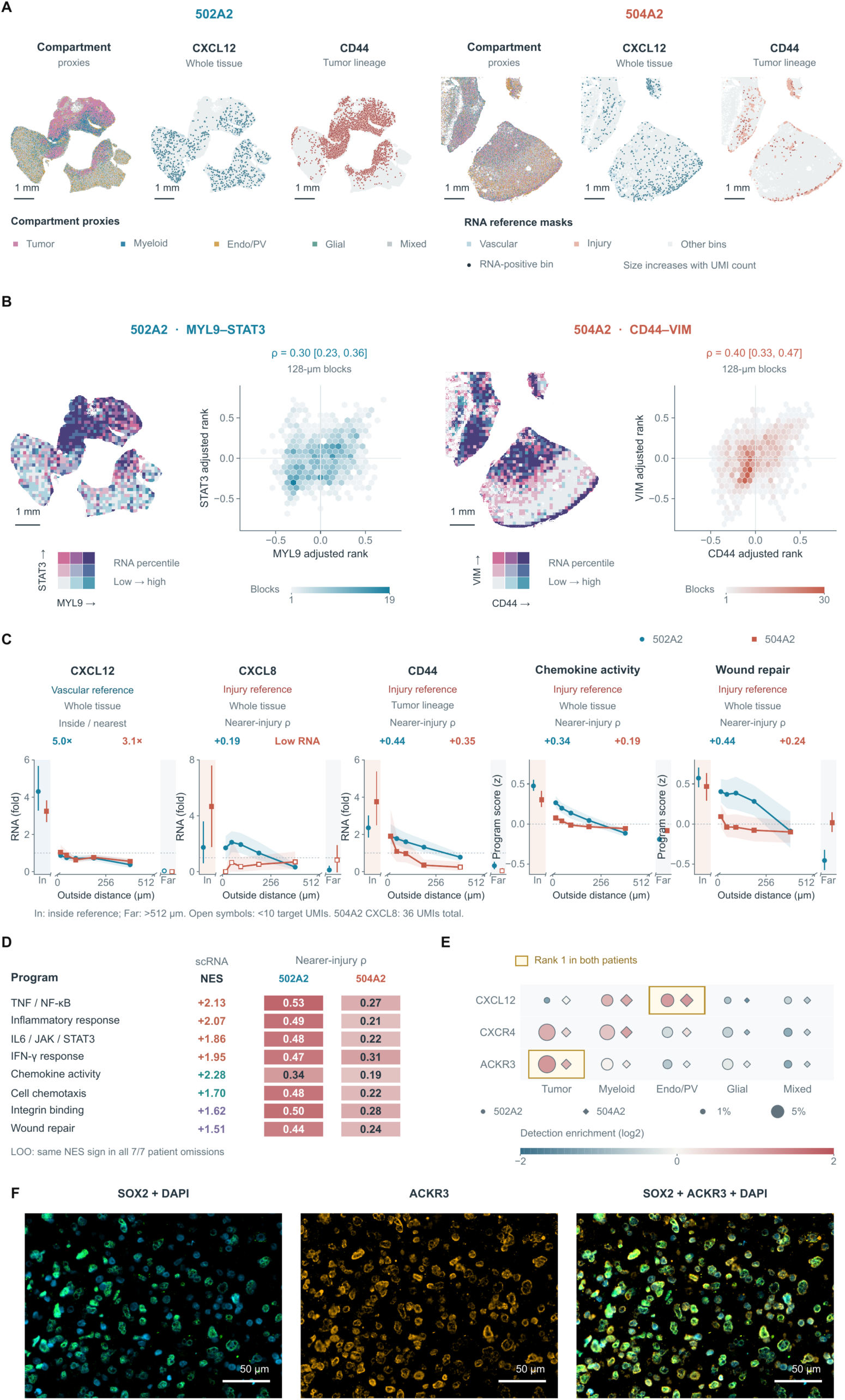
Spatial separation of vascular CXCL12 and injury-associated inflammatory–repair transcription. **A**, Native 16-µm-bin maps from TMZ-exposed sections 502A2 and 504A2, showing compartment proxies and CXCL12 in whole tissue or CD44 in tumor-lineage-enriched bins. **B**, MYL9–STAT3 and CD44–VIM maps with adjusted-rank hexagons at 128-µm resolution. Map colors indicate RNA percentile thirds and hexagon colors indicate block counts; annotations show correlations with conditional 95% spatial-bootstrap intervals. STAT3 RNA is not a measure of STAT3 phosphorylation. **C**, Relative RNA abundance or mean program z scores by distance from vascular- or injury-reference territories, with conditional 95% spatial-bootstrap intervals. Open symbols indicate fewer than 10 target unique molecular identifiers. Inside-reference and >512-µm categories are discrete groups; annotations show inside-to-nearest enrichment or adjusted external proximity correlations. **D**, Human single-cell GSEA NES alongside injury-proximity correlations for matched annotations; these are distinct statistics. **E**, Compartment-level RNA detection: symbol area denotes detection frequency and color log₂ detection enrichment; gold outlines identify the highest-ranked compartment in both sections. Scale bars in A,B, 1 mm. Intervals describe within-section uncertainty, not between-patient variability. **F**, Representative immunofluorescence images of the same tissue field showing SOX2 and DAPI (left), ACKR3 (middle), and merged SOX2, ACKR3, and DAPI (right). SOX2, green; ACKR3, gold; DAPI, cyan-blue. Scale bars, 50 µm.

CXCL12 abundance was approximately 5.0- and 3.1-fold higher inside vascular-reference territories than in the nearest external bands. By contrast, tumor-lineage-enriched CD44 increased toward injury-reference territories (adjusted proximity ρ = 0.44 and 0.35). Chemokine-activity and wound-repair scores showed corresponding injury-proximity associations (ρ = 0.34/0.19 and 0.44/0.24 across 502A2/504A2; **Fig. 2C,D**). CXCL8 showed a positive association in 502A2 (ρ = 0.19), whereas low detection limited interpretation in 504A2. Compartment-level detection placed CXCL12 preferentially in endothelial/perivascular bins and ACKR3 in tumor-enriched bins (**Fig. 2E**). Representative multiplex immunofluorescence additionally showed ACKR3 protein staining alongside SOX2 in the same tissue field (**Fig. 2F**). These observations distinguish vascular chemokine enrichment from injury-proximal inflammatory–repair transcription within the sampled sections.

Native-resolution maps localized CXCL12-positive vascular, CXCR4-positive myeloid and ACKR3-positive tumor-enriched bins in both sections (**Supplementary Fig. 3A**). Positive whole-tissue injury-proximity associations persisted across 64–256-µm blocks and top-5% to top-15% reference definitions in technical-covariate-adjusted sensitivity analyses (**Supplementary Fig. 3B,C**).

### Patient-resolved analyses nominate vascular–myeloid CXCL12–CXCR4 and vascular–tumor CXCL12–ACKR3 interactions

Cell-type expression profiles identified endothelial CXCL12, prominent myeloid CXCR4 and tumor-associated ACKR3 (**Fig. 3A**). CellChat supported endothelial-to-microglial and endothelial-to-macrophage CXCL12–CXCR4 interactions in all seven patients; LIANA supported these interactions in five and six patients, respectively. Direct endothelial-to-malignant CXCL12–CXCR4 predictions were less consistent (CellChat, 3/7; LIANA, 0/7), whereas CXCL12–ACKR3 was supported in five and three patients, respectively. Myeloid OSM- and TNF-associated predictions provided additional candidate routes to malignant cells (**Fig. 3B,C**).

**Figure 3.**
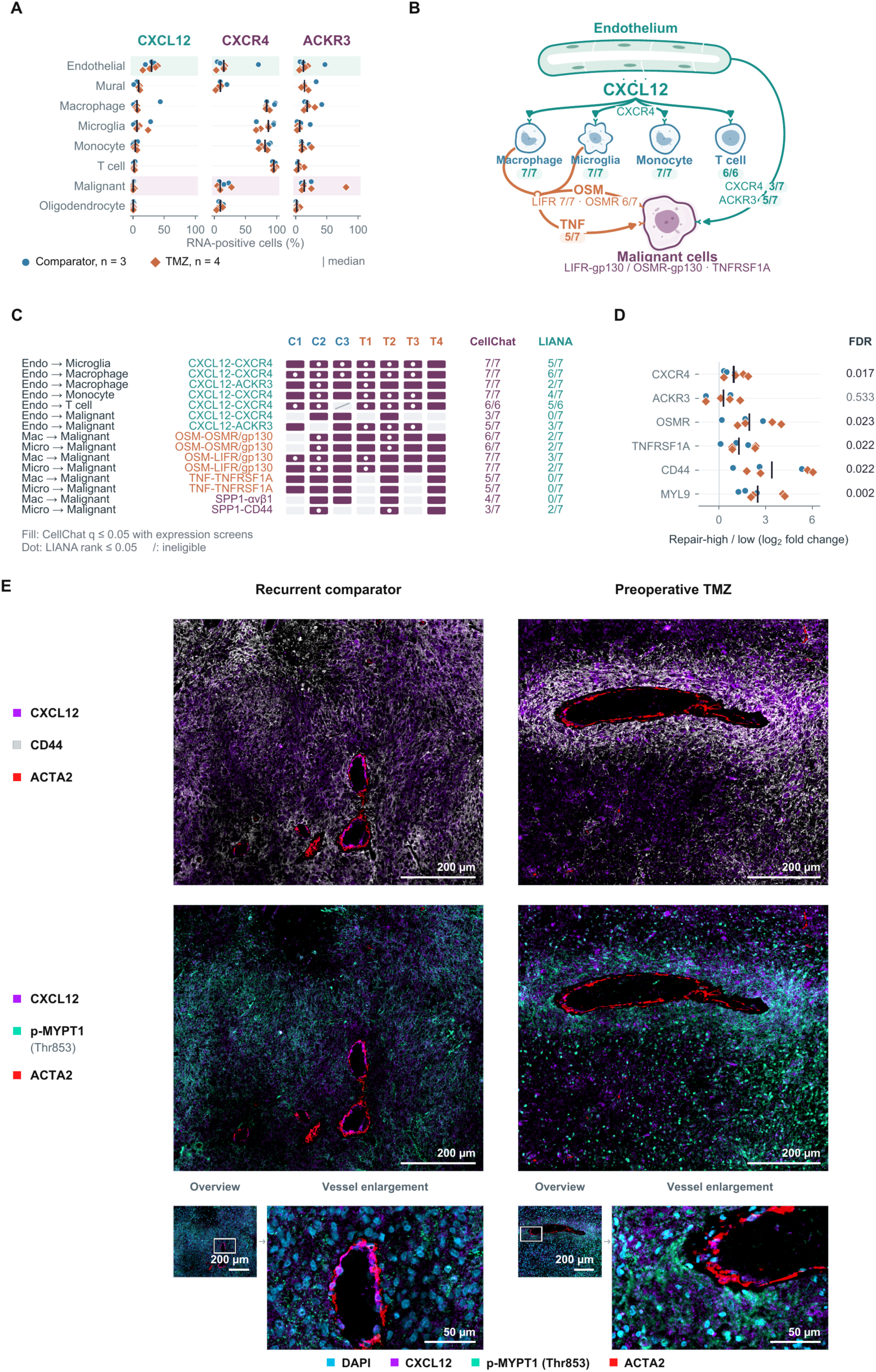
Candidate vascular–myeloid and vascular–glioblastoma chemokine interactions. **A**, RNA-positive fractions across eligible populations from three comparator and four preoperative-TMZ patients. Symbols represent patients and ticks indicate equal-patient medians. Communication analyses required at least 30 cells per population. **B**, Candidate ligand–receptor routes; fractions denote CellChat-supported patients among those with eligible populations, and arrows represent inferred rather than experimentally demonstrated interactions. **C**, Patient-level support: filled tiles indicate CellChat targeted q ≤ 0.05 with expression/standard screens; dots indicate LIANA specificity rank ≤ 0.05; slashes indicate ineligible populations. LIANA specificity ranks are not P values. Patient labels: C, recurrent comparator; T, preoperative TMZ. **D**, Repair-high versus repair-low malignant-cell expression. Symbols show paired patient log₂ fold changes, ticks show model estimates, and FDR values reflect adjustment within the corresponding joint testing family. **E**, Representative multiplex tissue images from a recurrent comparator without additional preoperative TMZ (left) and a preoperative-TMZ specimen (right). Top row, CXCL12, CD44 and ACTA2; middle row, CXCL12, p-MYPT1 (Thr853) and ACTA2; bottom row, DAPI, CXCL12, p-MYPT1 (Thr853) and ACTA2, shown as an overview and corresponding vessel enlargement for each specimen. Overview images show the same field across rows within each specimen; white boxes identify the regions shown in the enlargements. CXCL12, purple; CD44, white; ACTA2, red; p-MYPT1 (Thr853), green/teal; DAPI, cyan-blue. Scale bars, 200 µm in overviews and 50 µm in enlargements.

Repair-high malignant states expressed higher CXCR4, OSMR, TNFRSF1A, CD44 and MYL9 than repair-low states (FDR = 0.017, 0.023, 0.022, 0.022 and 0.002), without a significant ACKR3 difference (FDR = 0.533; **Fig. 3D**). Representative multiplex imaging showed qualitatively more prominent CD44 and p-MYPT1 signals in the preoperative-TMZ specimen than in the recurrent comparator (**Fig. 3E**). Matched views containing SOX2, Nestin, ACKR3/CXCR7 or p-MYPT1 provide further spatial context (**Supplementary Fig. 4A–D**). These analyses consistently nominate vascular–myeloid CXCL12–CXCR4 interactions and identify CXCL12–ACKR3 as a candidate vascular-to-glioblastoma interaction, alongside additional myeloid-derived signals associated with repair-high malignant states.

### MYL9 covaries with CXCL12, p-MYPT1 and nuclear p-STAT3 in human glioblastoma

Multiplex protein imaging provided an orthogonal assessment of these relationships (**Fig. 4A**). Across seven specimens, autofluorescence-corrected, locally adjusted MYL9 abundance correlated with CXCL12 (summary ρ = 0.61; 95% confidence interval [CI], 0.53–0.69), p-MYPT1 Thr853 (ρ = 0.48; 0.37–0.57) and nuclear p-STAT3 (ρ = 0.45; 0.37–0.53). Each association was positive in every specimen (**Fig. 4B**). Inflammatory, reactive/mesenchymal, cytoskeletal and invasion-associated protein scores also covaried (**Fig. 4C**).

**Figure 4.**
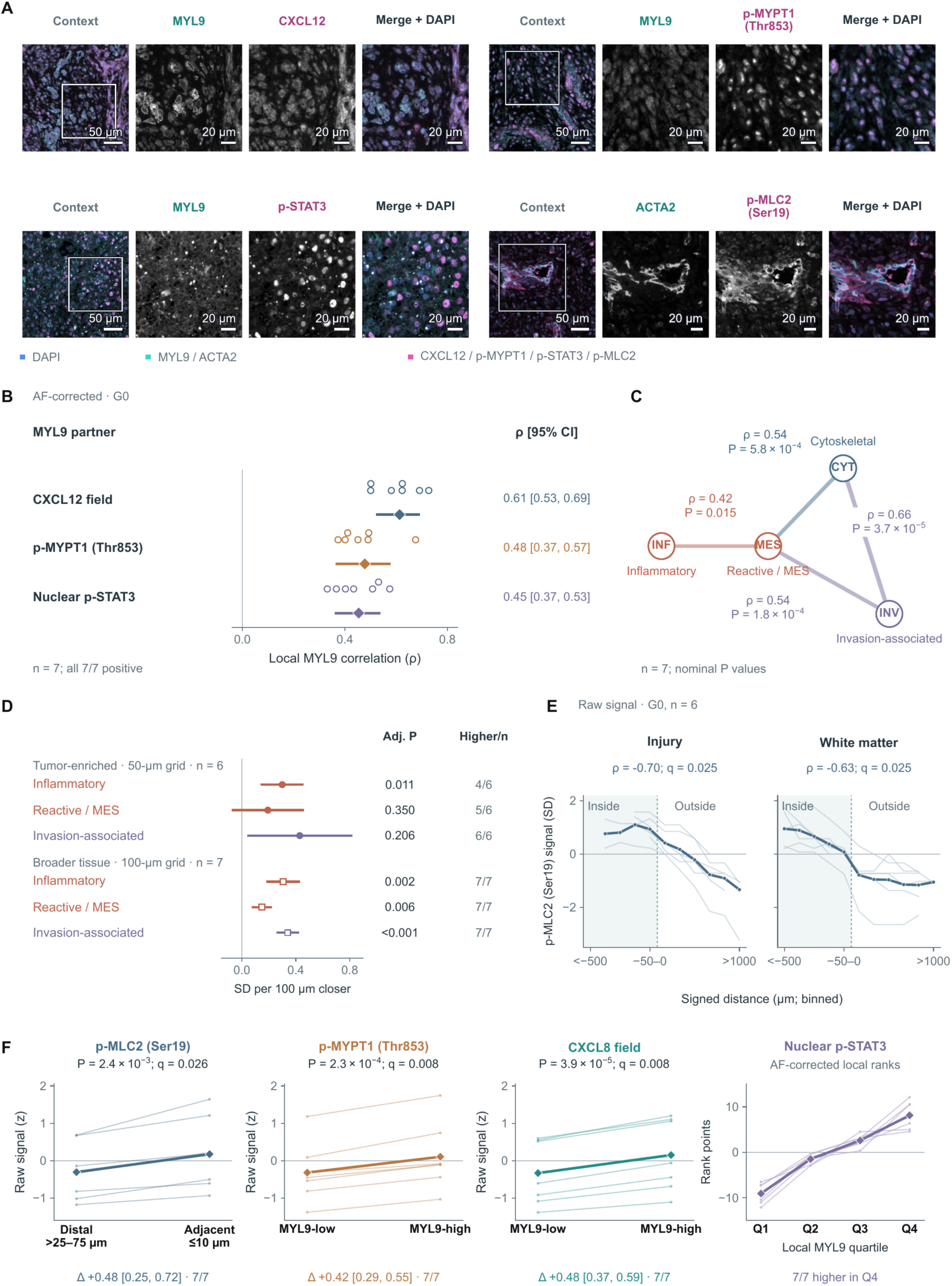
MYL9-associated chemokine and phosphoprotein states in human glioblastoma. **A**, Multiplex immunofluorescence fields. Upper-left, upper-right and lower-right groups are from NU0419; the lower-left group is from NU04849A1. Boxes indicate enlarged regions. DAPI, blue; MYL9 or ACTA2, cyan; indicated partner, magenta. Scale bars, 50 µm for overview images and 20 µm for enlargements. **B**, Autofluorescence-corrected, locally adjusted MYL9 correlations in G0-gated cells from seven specimens. Circles represent specimens; diamonds and bars show Fisher-z means and t-based 95% CIs. **C**, Protein-program correlations: inflammatory, reactive/mesenchymal, cytoskeletal (p-MLC2 and p-MYPT1), and invasion-associated (gelsolin, CXCR4 and ACKR3). Displayed P values are nominal. **D**, Injury-proximity effects in tumor-enriched tissue (50-µm grid; n = 6) and broader tissue (100-µm grid; n = 7), expressed as SD change per 100 µm closer to the reference territory. Bars show Hartung–Knapp–Sidik–Jonkman 95% CIs; adjusted P values reflect the corresponding prespecified testing families. **E**, Raw p-MLC2 Ser19 boundary profiles in G0 cells from six specimens; thin lines show individual specimens and thick lines show equal-specimen means. Within each specimen, Spearman correlations were calculated using 100-µm spatial-block medians and Fisher-z transformed. Two-sided one-sample t tests against zero were performed across specimens (df = 5). Displayed q values are Benjamini–Hochberg adjusted across 297 eligible marker-by-context tests in the G0/raw-signal/100-µm-block testing family. **F**, Vessel-distance and MYL9-state comparisons. The first three panels use raw G1 measurements from seven specimens. Δ denotes the equal-patient mean within-specimen difference with a t-based 95% CI; two-sided paired t tests were used. Displayed q values are Benjamini–Hochberg adjusted across 184 eligible vascular-distance tests for the p-MLC2 comparison and 1,436 eligible marker-state tests for the p-MYPT1 and CXCL8 comparisons. The fourth panel shows the autofluorescence-corrected G0 nuclear p-STAT3 profile across MYL9 quartiles and is descriptive; no quartile-comparison test is displayed. G0/G1 are analysis gates, not cell-cycle phases.

Within tumor-enriched regions, inflammation increased by 0.30 SD per 100 µm closer to injury-reference regions (n = 6 specimens; adjusted P = 0.011), whereas reactive/mesenchymal and invasion-associated effects did not meet the adjusted significance threshold (P = 0.350 and 0.206). All three programs increased toward injury-reference regions in the broader tissue analysis (n = 7; **Fig. 4D**). Raw p-MLC2 boundary profiles and vessel-distance/MYL9-state contrasts showed additional anatomical differences (**Fig. 4E,F**), but anatomical effects were attenuated after autofluorescence correction. The most consistent protein-level result was therefore local MYL9 covariation with CXCL12, p-MYPT1 and nuclear p-STAT3.

### TMZ promotes inflammatory–reactive reprogramming and enhanced glioblastoma migration

In PDX-derived glioblastoma cultures, TMZ exposure increased CD44, IRF9, and p75NTR immunofluorescence signals (**Fig. 5A,B**) and CXCL8, IL11 and GDF15 transcripts (**Fig. 5C**), recapitulating selected inflammatory and reactive features identified in the human analyses.

**Figure 5.**
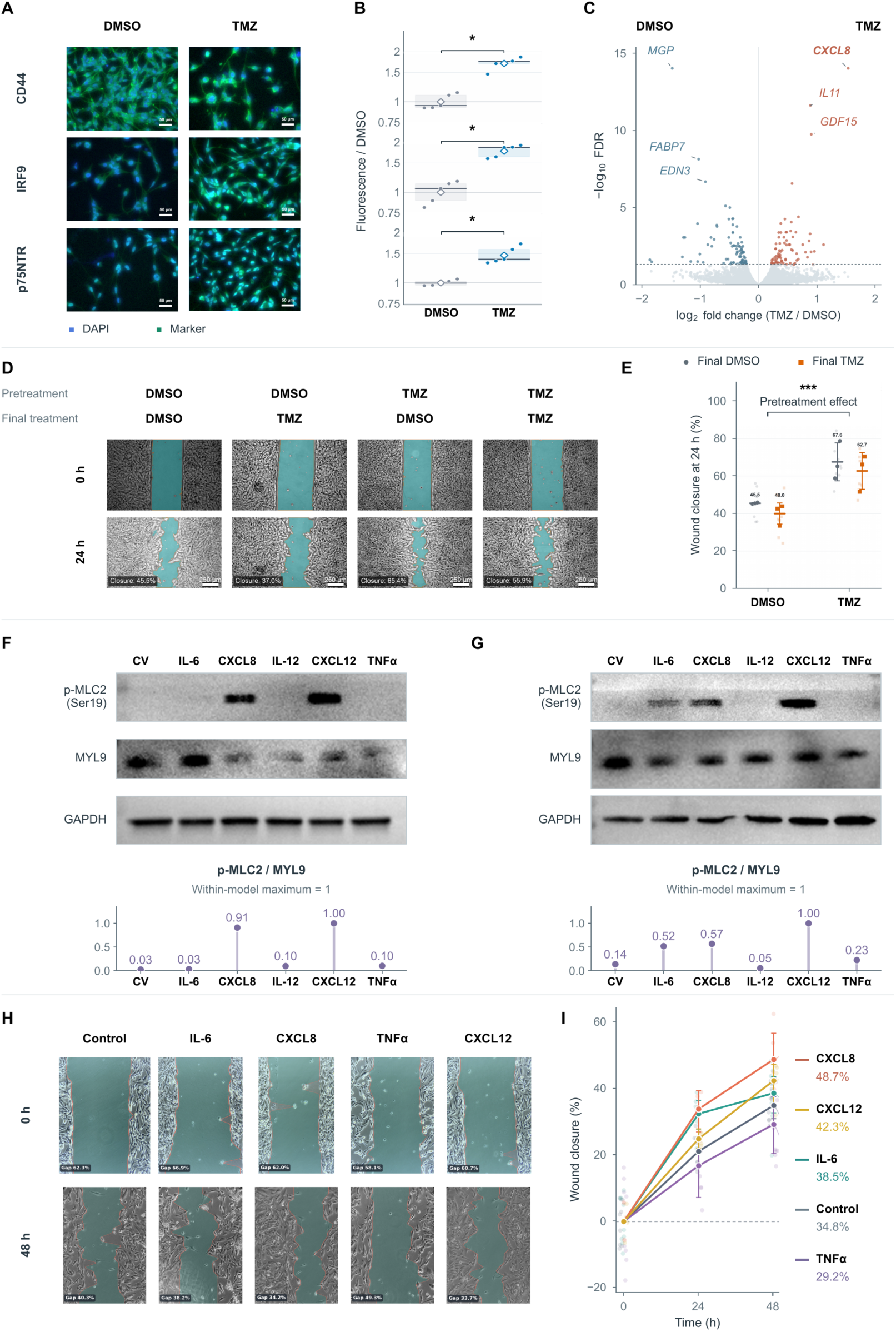
TMZ-associated reactive-state features and ligand-responsive scratch-closure and phosphoprotein readouts. **A**,**B**, Representative immunofluorescence images and corresponding quantification of CD44, IRF9 and p75NTR after TMZ or DMSO exposure in GBM43. Signals are shown relative to DMSO. Small symbols represent five technical image fields per condition; field-level measurements represent technical observations rather than independent biological replicates. DAPI, blue; indicated marker, green. Scale bars, 50 µm. **C**, Bulk RNA-sequencing volcano plot for TMZ versus DMSO; axes show log₂ fold change and −log₁₀ FDR, and the dashed line FDR = 0.05. Selected increased transcripts include CXCL8, IL11 and GDF15. **D**,**E**, Scratch assay after 48-h DMSO or TMZ pretreatment, followed by DMSO or TMZ for 24 h after scratch formation, with baseline and 24-h images and closure summaries. Mean closure was 45.5%, 40.0%, 67.6% and 62.7% for DMSO→DMSO, DMSO→TMZ, TMZ→DMSO and TMZ→TMZ, respectively. Image annotations denote individual fields rather than group means. Scale bars, 250 µm. **F**,**G**, Cytokine-stimulation immunoblots and displayed p-MLC2 Ser19/MYL9 densitometry in GBM38 and GBM43, respectively. Conditions are control vehicle (CV), IL-6, CXCL8, IL-12, CXCL12 and TNF-α. MYL9 and GAPDH bands are shown in both models; plotted ratios use MYL9 and are scaled to the maximum within each blot (maximum = 1). Values are descriptive within-blot measurements, not absolute phosphorylation stoichiometry. **H**,**I**, Representative cytokine scratch images and closure trajectories at 0, 24 and 48 hours. Mean 48-hour closure was 48.7% with CXCL8, 42.3% with CXCL12, 38.5% with IL-6, 29.2% with TNF-α and 34.8% in controls.

To test whether prior TMZ exposure altered subsequent scratch closure, GBM43 cells were pretreated with DMSO or TMZ for 48 h, scratched, and then maintained with DMSO or TMZ for 24 h. TMZ-pretreated cultures showed greater mean 24-hour closure than DMSO-pretreated cultures under either post-scratch treatment: 67.6% versus 45.5% with post-scratch DMSO and 62.7% versus 40.0% with post-scratch TMZ (**Fig. 5D,E**). Prior TMZ exposure was therefore associated with an approximately 22-percentage-point increase in closure under both conditions.

### CXCL8 and CXCL12 converge on actomyosin activation in glioblastoma cells

CXCL8 emerged from the TMZ-associated transcriptional changes, whereas CXCL12 emerged from human spatial analyses. We therefore tested these and other inflammatory ligands for their effects on cytoskeletal signaling. CXCL8 and CXCL12 induced robust p-MLC2 Ser19 signals in both PDX-derived cell lines (**Fig. 5F,G**). After normalizing the p-MLC2/MYL9 ratio to the maximum ratio within each blot, control/CXCL8/CXCL12 values were 0.03/0.91/1.00 and 0.14/0.57/1.00, respectively. IL-6 responses differed between models (0.03 and 0.52).

We next examined ligand-associated scratch closure. At 48 hours, mean closure was 48.7% with CXCL8, 42.3% with CXCL12, 38.5% with IL-6 and 29.2% with TNF-α, compared with 34.8% in control cultures (**Fig. 5H,I**). CXCL8 produced the largest increase in closure, whereas responses to the other ligands were smaller or directionally variable. Thus, CXCL8 and CXCL12 converged on a shared p-MLC2-associated cytoskeletal response, while their effects on scratch closure were ligand-dependent.

### Fasudil attenuates inflammatory signaling, actomyosin activation and glioblastoma migration

Because MYL9 covaried with ROCK-associated p-MYPT1 Thr853 in human tumors and CXCL8/CXCL12 increased p-MLC2 in glioblastoma cells, we next tested whether pharmacologic ROCK inhibition with fasudil would attenuate these inflammatory and cytoskeletal phenotypes. Bulk RNA profiling compared vehicle, TMZ and TMZ plus fasudil (n = 3 samples per condition; **Fig. 6A,B**). Relative to TMZ alone, the combination showed negative enrichment of TNF–NF-κB, inflammatory-response and IFN-γ-response programs (NES = −1.63, −1.65 and −2.33; q = 0.002, 0.002 and <0.001). The IL-6–JAK–STAT3 program did not meet q < 0.05 (NES = −1.36; q = 0.088).

**Figure 6.**
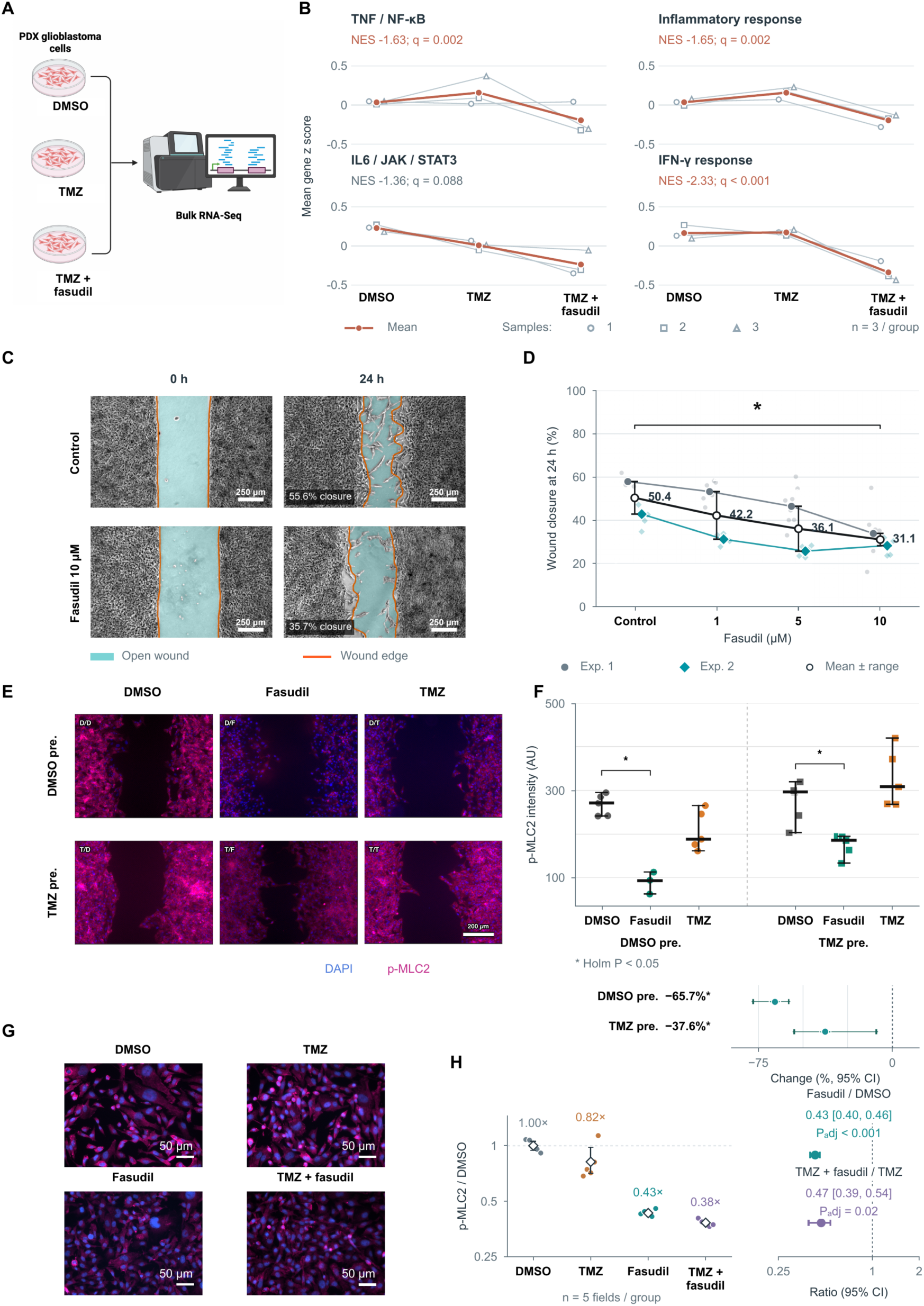
Fasudil attenuates inflammatory signaling, actomyosin activation and glioblastoma migration. **A**, Bulk RNA-sequencing design comparing DMSO, TMZ and TMZ plus fasudil. **B**, Sample-level program scores calculated as the mean of member-gene z scores derived from TMM log₂-CPM values standardized across all nine samples (n = 3 samples per condition). Symbols identify individual samples and bold lines connect condition means. Annotated NES and q values refer to preranked GSEA for TMZ plus fasudil versus TMZ, with Benjamini–Hochberg correction across the original 11-program family, including programs not displayed. These statistics are not tests of the plotted score means. IL-6–JAK–STAT3 did not meet q < 0.05. **C**,**D**, GBM38 scratch assay with control or 1, 5 or 10 µM fasudil. Two experimental trajectories and their mean with range are shown at 24 hours. Scale bars, 250 µm. **E**,**F**, p-MLC2/DAPI imaging and background-corrected quantification after DMSO or TMZ pretreatment followed by DMSO, fasudil or TMZ challenge. Contrast summaries show median percentage changes after fasudil challenge with 95% intervals; the panel specifies Holm-adjusted P < 0.05. Scale bars, 200 µm. **G**,**H**, GBM43 p-MLC2/DAPI images and quantification across DMSO, TMZ, fasudil and combination conditions. Five image fields per condition were analyzed. Contrast plots show fasudil-to-DMSO and combination-to-TMZ ratios with image-field bootstrap 95% intervals. These field-level intervals and statistical comparisons describe technical within-experiment variation and were not interpreted as evidence of independent biological replication. Scale bars, 50 µm.

Across two GBM38 scratch experiments, mean 24-hour closure decreased from 50.4% in control cultures to 42.2%, 36.1% and 31.1% with 1, 5 and 10 µM fasudil, respectively (**Fig. 6C,D**). Consistent with ROCK pathway inhibition, fasudil challenge also reduced median p-MLC2 signal by 65.7% after vehicle pretreatment and 37.6% after TMZ pretreatment (**Fig. 6E,F**). In a separate GBM43 imaging experiment, the combination-to-TMZ p-MLC2 ratio was 0.47 (95% image-field bootstrap interval, 0.39–0.54; five fields per condition; **Fig. 6G,H**). TMZ alone did not increase p-MLC2 above vehicle in that experiment. In an additional fasudil concentration series, p-MLC2 Ser19 and CD44 fluorescence signals were lower than control at all tested concentrations (1, 5 and 10 µM; **Supplementary Fig. 5A–C**). ROCK inhibition therefore attenuated inflammatory transcription and actomyosin activation and reduced glioblastoma migration in scratch assays.

### Orthotopic models recapitulate TMZ-associated inflammatory–repair programs and show prolonged survival with fasudil plus TMZ

Finally, we examined transcriptional responses and therapeutic outcomes in orthotopic glioblastoma models. Human-enriched single-cell tumor profiles from four independent mice per condition following TMZ or vehicle treatment were aggregated to one sample-level pseudobulk per mouse for differential-expression and pathway analyses (**Fig. 7A**). TMZ enriched hypoxia, TNF–NF-κB and inflammatory-response programs (NES = 1.78, 1.63 and 1.45; FDR < 0.001, <0.001 and 0.034, respectively), together with cell chemotaxis, integrin binding and wound healing (NES = 1.57, 1.59 and 1.30; FDR = 0.003, 0.005 and 0.034, respectively; **Fig. 7B**). Thus, orthotopic tumors reproduced several inflammatory, adhesion and tissue-repair features observed in human glioblastoma following TMZ exposure.

**Figure 7.**
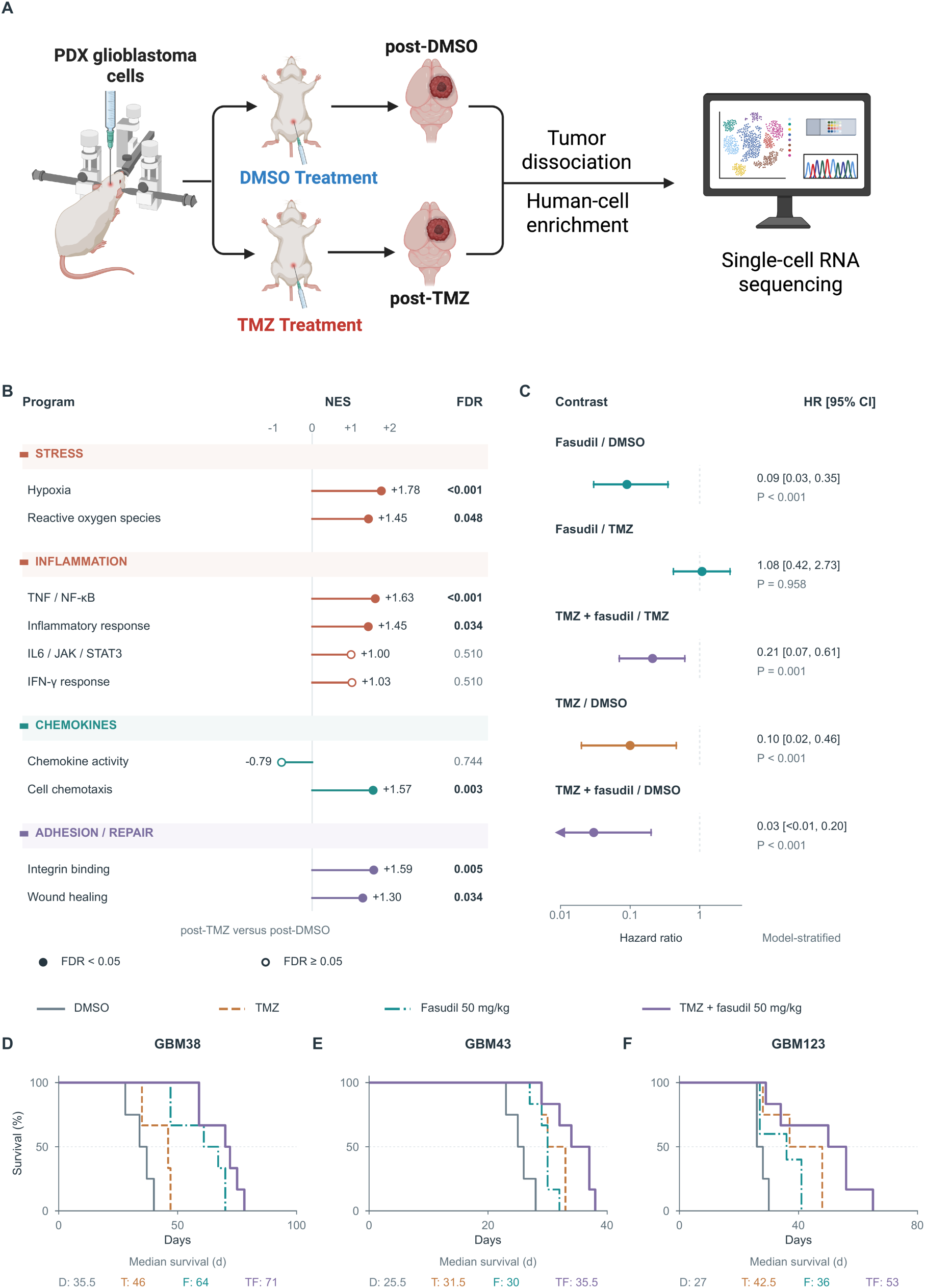
Orthotopic transcriptional responses to TMZ and survival with fasudil plus TMZ. **A**, Orthotopic implantation and transcriptional-profiling workflow: vehicle or TMZ exposure, tumor dissociation, human-cell enrichment and single-cell RNA sequencing. **B**, Targeted gene-set enrichment for the post-TMZ versus post-DMSO sample-level pseudobulk contrast. The displayed comparison comprised four independent mice per condition, with one pseudobulk library per mouse, representing 11,138 post-DMSO and 15,655 post-TMZ cells. Genes were ranked by sign(log₂ fold change) × −log₁₀(P value); enrichment used 20,000 permutations, with Benjamini–Hochberg correction across the ten displayed programs. Points denote NES; filled and open symbols denote FDR < 0.05 and FDR ≥ 0.05, respectively. **C**, Model-stratified Cox hazard ratios and 95% CIs for the indicated contrasts. A left-pointing arrow indicates an interval extending below the plotted range. **D**–**F**, Kaplan–Meier survival curves for GBM38, GBM43 and GBM123 treated with vehicle, TMZ, fasudil or TMZ plus fasudil. The displayed fasudil dose is 50 mg kg⁻¹. Median survival in vehicle/TMZ/fasudil/combination groups was 35.5/46/64/71 days for GBM38, 25.5/31.5/30/35.5 days for GBM43 and 27/42.5/36/53 days for GBM123. The combination had the longest median survival in each model. Hazard ratios and 95% confidence intervals were estimated using model-stratified Cox regression; P values were obtained from stratified log-rank tests.

We next evaluated vehicle, TMZ, fasudil, and the combination of fasudil plus TMZ across three PDX models (GBM38, GBM43, and GBM123). Across models, fasudil plus TMZ was associated with a lower hazard than TMZ alone (model-stratified Cox HR = 0.21; 95% CI, 0.07–0.61; stratified log-rank P = 0.001; **Fig. 7C**). Median survival was longer with the combination than with TMZ alone in all three models: 71 versus 46 days in GBM38, 35.5 versus 31.5 days in GBM43 and 53 versus 42.5 days in GBM123 (**Fig. 7D–F**). The combination produced the longest median survival in each model despite model-dependent differences in monotherapy responses. These orthotopic experiments recapitulated several components of the TMZ-associated inflammatory–repair phenotype and showed prolonged survival with combined ROCK inhibition and TMZ.

## DISCUSSION

A key feature of this study is the analysis of recurrent human glioblastoma tissue obtained following a defined 5-day presurgical TMZ exposure, providing a direct view of treatment-associated remodeling in the intact tumor microenvironment. This study identifies coordinated inflammatory and tissue-repair programs in recurrent human glioblastoma and places them within spatially structured chemokine environments linked to cytoskeletal phenotypes.

Across patients, inflammatory transcription covaried with adhesion and wound-repair programs, while MYL9 consistently associated with local CXCL12, p-MYPT1 and nuclear p-STAT3 protein signals. Controlled TMZ exposure reproduced selected inflammatory and reactive features and increased subsequent scratch closure, whereas CXCL8 and CXCL12 increased MLC2 phosphorylation. ROCK inhibition with fasudil attenuated inflammatory programs, migration and selected actomyosin readouts, and fasudil plus TMZ prolonged survival relative to TMZ alone across three orthotopic models. Together, these findings support a treatment-associated inflammatory–repair state coupled to cytoskeletal remodeling and identify ROCK inhibition as a pharmacologically tractable therapeutic strategy.

The human data should be interpreted in the context of recurrent, previously treated disease. Both groups had received prior TMZ and radiotherapy; the comparison therefore addresses additional preoperative TMZ rather than treatment-naïve versus TMZ-exposed tumors. The small observational cohort cannot distinguish treatment-induced reprogramming from selection of pre-existing states or other patient and microenvironmental differences. Nevertheless, the direction of the transcriptional differences was preserved across leave-one-patient-out analyses, and, independently of treatment group, inflammatory programs covaried with integrin-binding and wound-healing transcription within every tumor. These observations support coordinated inflammatory–repair biology while stopping short of establishing a treatment-induced state transition. Therapy-induced senescence could contribute to this phenotype, given prior evidence linking TMZ to NF-κB-associated senescence and secretory programs in glioblastoma [17,18], but senescence was not directly measured here.

The spatial data further suggest that this biology is multicompartmental rather than exclusively tumor-cell autonomous. CXCL12 localized preferentially to vascular territories, and patient-resolved communication analyses consistently nominated vascular–myeloid CXCL12–CXCR4 interactions. In contrast, CXCL12–ACKR3 represented a candidate vascular-to-glioblastoma route, alongside myeloid OSM- and TNF-associated signaling to malignant cells. Previous work similarly identified heterogeneous ACKR3 expression in glioblastoma tissue, with limited direct effects on proliferation or invasion in tested culture models, supporting a context-dependent role for ACKR3 in glioblastoma [19]. These findings are compatible with established roles for inflammatory cytokines and chemokines in glioblastoma state regulation [4,20], but spatial proximity and computationally inferred communication do not establish ligand–receptor signaling. Notably, the human CXCL12 observations and experimental CXCL8 effects nominate complementary inflammatory inputs rather than a single linear receptor pathway.

Protein imaging connected these inflammatory environments to cytoskeletal regulation. MYL9 abundance correlated with CXCL12, ROCK-associated p-MYPT1 and nuclear p-STAT3 across all seven specimens, providing orthogonal evidence for coordinated chemokine, inflammatory and actomyosin-associated states. However, co-abundance does not establish pathway order or kinase dependence, and some anatomical gradients were sensitive to autofluorescence correction. Injury-proximal inflammation was more robust than corresponding reactive/mesenchymal or invasion-associated gradients, emphasizing that injury, hypoxia and tumor growth may represent overlapping rather than separable spatial processes [7,8,10,11].

In vitro, TMZ increased CXCL8 and selected reactive-state markers and was associated with greater subsequent scratch closure. CXCL8 and CXCL12 induced prominent p-MLC2 responses in two glioblastoma models, while CXCL8 produced the largest increase in scratch closure. These findings are consistent with prior reports of TMZ-associated plasticity and CXCL8-linked glioma invasion [14,21], but they do not establish that chemokine-induced p-MLC2 directly mediates migration. Receptor perturbation, live-cell migration measurements and rescue experiments will be required to determine which receptors and downstream signals mediate these effects.

Pharmacological ROCK inhibition provided an additional test of this framework. Fasudil reduced TNF–NF-κB, inflammatory-response and IFN-γ-response programs and decreased scratch closure, with accompanying p-MLC2 reduction in selected experimental contexts. These effects do not establish ROCK isoform-specific dependence, and ROCK1 and ROCK2 may exert context-dependent effects in glioblastoma [22]. More importantly, fasudil plus TMZ prolonged survival relative to TMZ alone across all three orthotopic models, with a model-stratified hazard ratio of 0.21. Prior studies have described fasudil anti-invasive activity and fasudil–TMZ sensitization [15,16]; the present study extends this work by combining human inflammatory, spatial and cytoskeletal profiling with experimental evaluation of fasudil. The survival benefit does not establish synergy or prove that suppression of invasion is the operative mechanism, as effects on tumor growth, treatment sensitivity or the host microenvironment remain possible.

Several limitations remain, including the small human cohort, two spatial RNA specimens, inferred malignant and tumor-enriched compartments, and limited independent biological replication for selected experimental assays.

Together, these findings link treatment-associated inflammatory–repair programs to cytoskeletal remodeling in glioblastoma. Survival prolongation across three orthotopic models supports further translational development of fasudil plus TMZ for glioblastoma.

## METHODS

### Human specimens and study design

The human cohort comprised seven patients with recurrent glioblastoma. Four patients were enrolled in the phase 0 TMZ-only presurgical subgroup (Sub-group S1) of NCT05236036 and received temozolomide (TMZ; 150 mg/m² orally once daily for 5 consecutive days) before surgical resection. Three recurrent comparator patients underwent resection without additional preoperative TMZ. All seven patients had previously received radiotherapy and TMZ as part of standard-of-care treatment; thus, the primary comparison evaluated the association of additional preoperative TMZ exposure rather than treatment-naïve versus TMZ-exposed disease. Samples from patients receiving preoperative TMZ plus mycophenolate mofetil (MMF) were excluded from the present analysis.

Human analyses included malignant-cell single-cell RNA profiles, spatial RNA profiling of sections 502A2 and 504A2, and multiplex protein imaging across seven specimens. The spatial RNA sections were derived from preoperative-TMZ, IDH-wildtype glioblastomas. Different analytical modalities were not treated as independent patients or biological replicates.

Human tumor specimens were obtained under protocols approved by the Northwestern University Institutional Review Board (STU00215766-MOD0066). All participants provided written informed consent. All procedures involving human specimens were conducted in accordance with institutional guidelines and the Declaration of Helsinki.

### Animal studies and orthotopic xenografts

All animal procedures were approved by the Northwestern University Institutional Animal Care and Use Committee (IS00021383) and were performed in accordance with institutional guidelines. Male and female athymic nude mice (nu/nu; Charles River Laboratories) were maintained under standard temperature- and humidity-controlled conditions with a 12-h light/dark cycle and ad libitum access to food and water. Animals were anesthetized with ketamine/xylazine for intracranial implantation and received perioperative analgesia according to institutional protocols.

GBM38, GBM43 and GBM123 patient-derived glioblastoma cells were stereotactically implanted into the right cerebral hemisphere through a cranial burr hole. 1.5 × 10^5^ cells were implanted per mouse to a depth of approximately 3 mm from the dura. After 7 days of tumor establishment, mice were assigned to vehicle, TMZ, fasudil, or TMZ plus fasudil treatment. TMZ (2.5 mg kg⁻¹) and fasudil (50 mg kg⁻¹) were administered intraperitoneally once daily for 10 consecutive days; mice in the combination group received both agents on the same schedule. TMZ was prepared in 5% DMSO/95% PBS, and fasudil was prepared in PBS.

Animals were monitored for clinical evidence of tumor progression and euthanized at prespecified humane endpoints, including neurological deterioration, substantial weight loss, or moribund condition. Survival was measured from intracranial implantation to euthanasia at the humane endpoint. Group sizes were GBM38: vehicle n = 4, TMZ n = 3, fasudil n = 6, combination n = 6; GBM43: vehicle n = 4, TMZ n = 4, fasudil n = 6, combination n = 6; GBM123: vehicle n = 4, TMZ n = 4, fasudil n = 5, combination n = 6.

### Single-cell RNA sequencing

Cell number and viability were assessed using a Nexcelom Cellometer Auto2000 with acridine orange/propidium iodide staining. Approximately 16,000 cells were loaded onto the Chromium X Controller (10x Genomics) using a Chromium GEM-X Single Cell 3′ Chip Kit v4. cDNA and sequencing libraries were generated using the GEM-X Single Cell 3′ Kit v4 and Dual Index Kit TT Set A according to the manufacturer’s protocol. Library quality and concentration were assessed using the Agilent Bioanalyzer High Sensitivity DNA Kit and Qubit DNA High Sensitivity assay, respectively. Multiplexed libraries were pooled and sequenced on an Illumina NovaSeq X Plus using 28-bp Read 1 for cell barcode and unique molecular identifier and 90-bp Read 2 for transcript sequence. Single-cell library preparation and sequencing were performed at the Northwestern University NUseq Core.

### Single-cell RNA processing and annotation

The source single-cell dataset contained 269,297 cells and 38,606 features across 16 libraries. Analyses reported here used 165,625 cells from three recurrent comparator and four preoperative-TMZ specimens; cells from the TMZ-plus-MMF condition were excluded. Integer counts were retained as the primary expression layer. Counts were normalized to 10,000 counts per cell and log-transformed as log1p(CP10K), with duplicate gene symbols summed before normalization.

The original integrated expression embedding and source clustering were preserved; no additional scVI integration, Harmony correction or UMAP refitting was performed for the final annotation analysis. Broad-lineage and Level-2 identities were assigned using CellTypist predictions [23] from immune, brain and vascular reference models together with predefined multigene marker panels, local-neighborhood support and model-agreement criteria. Broad lineages comprised neural/glial, myeloid, lymphoid, vascular/stromal, other and unresolved populations. Level-2 annotations included oligodendrocytes, oligodendrocyte precursor cells, astrocytic cells, neural-progenitor-like cells, microglia, macrophages, monocytes, T cells, NK cells, B cells, endothelial cells, fibroblasts, pericytes and smooth-muscle cells. Cells with insufficient or conflicting evidence were retained as unresolved rather than forcibly assigned. Existing doublet annotations and review flags were preserved; putative doublets were not globally excluded.

### RNA-inferred copy-number analysis and malignant-cell selection

RNA-inferred copy-number profiles were generated independently of expression-based annotation using a custom chromosome-smoothing procedure across 14,575 genome-ordered genes. Within each chromosome, expression was smoothed using 101-gene windows advanced in 10-gene steps. Calibration used expression-supported immune, vascular and putative oligodendrocyte reference populations, with separate fitting, calibration and evaluation subsets. Treatment condition and pre-existing malignancy calls were not used for CNV classification.

Chromosome 7 gain and chromosome 10 loss were evaluated independently, with a positive directional call requiring at least 60% of informative windows to support the expected direction at the prespecified 95th-percentile calibration threshold. Cells lacking sufficient information were retained as low-information rather than forced into positive or negative categories. RNA-CNV calls were therefore treated as RNA-expression signatures rather than DNA-confirmed copy-number alterations. Within the seven-patient comparator-plus-TMZ cohort, 32,991 cells meeting the joint chromosome 7 gain/chromosome 10 loss criterion were retained for malignant-cell analyses. Copy-number profiles were summarized across 535 genomic bins for visualization, and this pattern was considered part of the selection procedure rather than independent validation of malignant identity.

### Transcriptional programs and patient-resolved analyses

Raw integer RNA counts from malignant cells meeting the joint chromosome 7 gain/chromosome 10 loss criterion were summed within each patient, combining specimens from the same patient before analysis. The analysis included seven patient-level pseudobulks: three recurrent comparators and four preoperative-TMZ patients, comprising 32,991 malignant cells. All patients exceeded the minimum of 100 malignant cells, and TMZ-plus-MMF samples were excluded. Low-expression features were removed using edgeR filterByExpr with the treatment design, retaining 19,582 features, followed by trimmed mean of M-values (TMM) normalization. For pathway ranking, normalized counts were analyzed using limma–voom [24,25], with an unpaired design, ∼ condition, with recurrent comparator as the reference, followed by robust empirical-Bayes moderation. No additional covariates were included. Genes were ranked by the signed moderated t statistic for preoperative TMZ versus recurrent comparator, without a significance or fold-change threshold. Duplicate gene symbols were represented by the feature with the greatest summed raw count across patients, yielding 19,577 ranked genes. Patients, rather than individual cells or specimens, were the biological replicates. Preranked gene-set enrichment analysis [26] used human MSigDB v2026.1.Hs Hallmark, Gene Ontology biological process, cellular component and molecular function, and C2:CP:PID collections. Gene sets were required to contain 15–500 measured genes and at least 20% of their original membership. Enrichment was computed using fgsea v1.30.0 fgseaMultilevel with gseaParam = 1, scoreType = “std”, sampleSize = 101, nPermSimple = 10,000, eps = 1e-10 and seed 20260907. Benjamini–Hochberg correction was applied separately across all 50 eligible Hallmark sets, all 4,973 eligible Gene Ontology sets pooled across the three domains, and all 180 eligible PID sets. Selected pathways retained their full-family adjusted q values. Seven leave-one-patient-out analyses refitted TMM normalization, voom and robust empirical-Bayes models while retaining the full-data gene universe and representative-symbol mapping. Enrichment was recalculated using fgseaSimple with 1,000 permutations per omission; enrichment-score signs were used to assess directional stability. These analyses assessed patient influence rather than independent replication.

Within-patient inflammation–integrin-binding and inflammation–wound-healing relationships used disjoint signatures with adjustment for technical covariates. Pair-specific inflammation deciles were summarized using equal-patient medians and interquartile ranges, and seven patient-specific partial Spearman correlations were summarized for each comparison. Repair-high versus repair-low malignant-cell contrasts were represented as paired patient log₂ fold changes, with model estimates and FDR values adjusted within the corresponding joint testing family.

### Candidate cell–cell communication

Candidate ligand–receptor interactions were evaluated separately for eligible patient cell populations using CellChat and LIANA [27,28]. Eligibility required at least 30 cells per population. CellChat support required the reported targeted Benjamini–Hochberg q ≤ 0.05 criterion together with expression and standard screening criteria. LIANA support used specificity rank ≤ 0.05; this rank was not interpreted as a P value. Interaction summaries reported supported patients among eligible populations, with ineligible populations distinguished from eligible but unsupported comparisons.

### Spatial transcriptomics acquisition and preprocessing

Spatial transcriptomic data were generated using the 10x Genomics Visium HD H1 platform with HD probe-based v1 chemistry and CytAssist imaging. Raw sequencing data were processed with Space Ranger count v4.0.1 using the Visium Human Transcriptome Probe Set v2.1.0 and GRCh38-2024-A reference. Tissue images were registered using the recorded Loupe fiducial-alignment files.

Analyses used the Space Ranger filtered 16-µm square-binned matrices rather than segmented-cell matrices. Bins were retained when located within the Space Ranger tissue footprint, contained at least 50 detected features, had mitochondrial transcript content ≤25%, and exceeded sample-specific lower outlier thresholds defined as median − 4.5 × median absolute deviation for total counts and detected features. High-count and high-feature bins were not excluded solely as upper outliers. After quality control, 62,284 bins from section 502A2 and 73,614 bins from section 504A2 were retained. Primary spatial analyses used these 16-µm bins directly without spatial interpolation, convolution or score smoothing.

### Spatial normalization, program scoring and compartment definition

Expression within each spatial bin was normalized to 10,000 total counts and log-transformed as log1p(CP10K). Gene-set scores were calculated as the unweighted mean of normalized expression across measured genes in each predefined program. Scores were standardized within section for visualization and converted to empirical within-section percentiles; these percentiles represent relative spatial rank within each section rather than between-section fold changes.

For primary injury-proximity analyses, genes shared between the injury-reference signature and the tested transcriptional program were removed before score reconstruction. Scores were subsequently adjusted within section for transcript depth, detected-feature count and transcriptomic compartment proxies.

Malignant-like, myeloid-rich, vascular/perivascular and astrocytic/CNS-like compartment scores were generated from predefined marker panels. Each bin was assigned to the highest-scoring compartment when the leading standardized score was non-negative and exceeded the second-highest score by at least 0.03; otherwise, it was classified as mixed/low-confidence. Tumor-lineage-enriched bins comprised bins at or above the section-specific 75th percentile of the malignant-like score and were interpreted as transcriptomic proxies rather than histological or DNA-CNV-confirmed tumor calls. CXCL12 and CXCL8 analyses used whole tissue, whereas CD44 proximity analyses used tumor-lineage-enriched bins.

RNA-defined injury-reference regions were generated independently of the tested inflammatory, chemokine, chemotaxis, integrin-binding and wound-healing programs. After removal of overlapping genes, the reference comprised 37 measured genes. Adjusted reference scores were summarized in fixed 8 × 8-bin blocks (128 × 128 µm); blocks were retained when they contained at least 16 tissue bins, fell within the upper 20% of the adjusted reference-score distribution and belonged to an 8-connected component containing at least three selected blocks. Supplementary sensitivity analyses used alternative top-5%, top-10% and top-15% reference definitions and block sizes of 64–256 µm, as specified in Supplementary Fig. 3. These masks represent computational transcriptomic reference regions rather than pathologist-defined injury boundaries. Reference-distance profiles were summarized using conditional 95% spatial-bootstrap intervals.

### Multiplex immunofluorescence acquisition and image processing

Formalin-fixed tissue sections were analyzed using the Lunaphore COMET multiplex-immunofluorescence platform. Acquisition comprised an initial DAPI and unstained-autofluorescence reference cycle followed by 20 two-marker staining cycles, producing 45-channel OME-BigTIFF images at a native pixel size of 0.28 µm. Native-resolution images were used for segmentation and marker quantification.

Image registration was assessed using acquisition metadata and native-resolution overlay quality control. Nuclear segmentation was performed on DAPI images using a locked fluorescence-nuclei instance-segmentation model selected using held-out expert annotations spanning diverse tissue habitats. Marker measurements were quantified within the nucleus, a 1–3-µm perinuclear ring and a bounded cell proxy extending up to 4 µm from the nuclear boundary; the latter was generated using bounded Voronoi assignment and was not interpreted as a directly observed whole-cell boundary.

For each compartment, intensity summaries included mean, median and 90th-percentile signal together with pixel counts and saturation metrics. Autofluorescence correction was performed separately for TRITC and Cy5 channels using corresponding unstained reference channels acquired during the initial cycle. Raw measurements were retained, with autofluorescence-corrected values used only where prespecified. Eligible nuclei had an area of 12–400 µm², a finite uncensored DAPI measurement and no contact with a tile-read boundary; subsequent tissue and pathology-validity masks could only make eligibility more restrictive.

### Multiplex cell populations and spatial reference regions

G0 and G1 denote analysis gates rather than cell-cycle phases. The strict G0 tumor-enriched glial gate required positive evidence for at least two of SOX2, OLIG2, EGFR and PDGFRA together with absence of CD3, IBA1, CD68, CD31 and ACTA2 evidence. CD44, Nestin, RRM2 and Ki-67 were not used to define admission to this gate. G0 cells were therefore panel-enriched glial/tumor-associated cells rather than genetically confirmed malignant cells. The broader G1 population corresponded to the prespecified GBM-enriched analysis gate and was used for the principal spatial analyses, with stricter gates retained for sensitivity analyses.

White-matter reference regions were defined using a pathologist-reviewed MBP-positive tract mask and intersected with tissue- and pathology-valid regions. Cells within the mask or a prespecified 100-µm peritract buffer were considered white-matter-associated. Injury-reference regions were defined only within white matter using three independent spatial-context features: high APP signal, high SMI32 signal and MBP disruption or local depletion. After robust standardization within white matter, prespecified thresholds were z ≥ 2.0 for APP, z ≥ 1.5 for SMI32 and z ≥ 1.0 for MBP disruption. The primary injury-reference mask required at least two of the three concordant features; morphological opening and closing were applied and components <400 µm² were removed. APP and SMI32 were interpreted as high-intensity spatial-context measurements rather than definitive morphological calls. Signed Euclidean distance fields were calculated from the reference masks, with injury proximity analyzed over 0–250 µm.

MYL9 partner analyses used autofluorescence-corrected, locally adjusted ranks and specimen-level correlations in G0 cells (n = 7), summarized using Fisher-z means and t-based 95% confidence intervals. Injury-proximity effects were evaluated in tumor-enriched tissue (n = 6) and broader tissue (n = 7) using 50-µm and 100-µm grids, respectively, with Hartung–Knapp–Sidik–Jonkman confidence intervals and multiplicity-adjusted P values. Raw p-MLC2 boundary profiles used six specimens. For vessel-distance and MYL9-state analyses, the first three comparisons used raw G1 measurements, whereas the nuclear p-STAT3 comparison used autofluorescence-corrected G0 measurements (n = 7 each).

### Glioblastoma cultures and treatment exposures

Patient-derived GBM38 and GBM43 glioblastoma cells were used for in vitro experiments. Temozolomide (TMZ) was administered at 50 µM for 48 h, with DMSO-treated cultures serving as vehicle controls. TMZ-versus-vehicle RNA and immunofluorescence assays assessed cytokine-associated transcripts and CD44, IRF9 and p75NTR signals. Fasudil scratch-closure experiments were performed in GBM38, and phospho-MLC2 imaging experiments were performed in GBM38 and GBM43.

### Bulk RNA sequencing, differential expression and gene-set enrichment

GBM43 cells were analyzed using three biological samples per condition for DMSO, TMZ, and TMZ plus fasudil. Gene-level counts were generated from aligned BAM files using Rsubread featureCounts and the Ensembl GRCh38 release 114 annotation. Reads had been aligned using STAR v2.7.11b to a GRCh38/Ensembl release 114 reference including ERCC sequences.

Exonic reads were summarized by gene_id using unstranded, single-end counting, excluding multimapping reads and reads assigned to multiple features. Following filterByExpr filtering and TMM normalization, 14,984 genes were retained. Differential expression was assessed using robust edgeR quasi-likelihood generalized linear models including treatment condition and replicate block. Contrasts were extracted from the common fitted model, and gene-level P values were adjusted using the Benjamini–Hochberg procedure.

For gene-set enrichment, genes were ranked by sign(log₂ fold change) × √F, where F was the edgeR quasi-likelihood test statistic from the common nine-sample model, ∼ 0 + condition + replicate_block. Duplicate gene symbols were represented by the feature with the greatest absolute ranking statistic, yielding 14,976 unique ranked symbols. The targeted analysis used frozen gene memberships for ten programs matched to the human analysis—six Hallmark and four Gene Ontology programs—together with an 82-gene injury-reference signature. Enrichment was performed using fgsea v1.30.0 fgseaMultilevel with minSize = 10, maxSize = 5,000, gseaParam = 1, scoreType = “std”, sampleSize = 101, nPermSimple = 20,000 and eps = 1e-10. The seed for the TMZ-plus-fasudil versus TMZ contrast displayed in Fig. 6 was 20260912. Benjamini–Hochberg correction was applied across the original 11-program family separately within each contrast, including the injury-reference signature, although Fig. 6 displays only four inflammatory programs. For visualization, nonconstant member-gene TMM log₂-CPM values were standardized across all nine samples using their gene-wise mean and sample standard deviation, and program scores were calculated as the unweighted mean of member-gene z scores. Annotated NES and q values derive from the ranked-gene enrichment analysis and are not tests of the displayed program-score means.

### Scratch-closure assays

For the TMZ pretreatment experiment, cells were exposed to DMSO or TMZ (50 µM) for 48 h before scratch formation, followed by terminal treatment with DMSO or TMZ (50 µM). Scratch closure was quantified from baseline to 24 h after scratch formation. Cytokine-stimulation experiments compared control with IL-6 (50 ng/mL), CXCL8 (100 ng/mL), CXCL12 (100 ng/mL), and TNF-α (50 ng/mL), with closure assessed at 0, 24 and 48 h. The GBM38 fasudil concentration series tested 1, 5 and 10 µM fasudil in two independent experiments, with 24-h closure summarized using the mean and range.

### Cytokine stimulation and immunoblotting

For cytokine-stimulation immunoblotting, GBM38 and GBM43 cells were exposed for 30 min to vehicle control, IL-6 (50 ng/mL), CXCL8 (100 ng/mL), IL-12 (20 ng/mL), CXCL12 (100 ng/mL), or TNF-α (50 ng/mL). Phospho-MLC2 Ser19 was assessed by immunoblotting, with densitometry calculated as background-corrected phospho-MLC2 Ser19 normalized to total MYL9. GAPDH was shown as a loading-control band but was not used as the densitometric denominator.

For TMZ/fasudil phosphoprotein experiments, TMZ was administered at 50 µM for 48 h and fasudil at 20 µM for 48 h. Four-condition imaging compared DMSO, TMZ, fasudil and TMZ plus fasudil. Individual image fields were treated as technical observations rather than independent biological replicates. In the separate fasudil concentration-series experiment, GBM43 cells were treated with vehicle or fasudil (1, 5 or 10 µM) for 24 h and analyzed by immunofluorescence for p-MLC2 Ser19 and CD44.

### Antibodies and immunodetection

Primary antibodies used for multiplex immunofluorescence included phospho-MYPT1 Thr853 (Thermo Fisher Scientific, PA5-40248), phospho-MLC2 Ser19 (Cell Signaling Technology, 3675S), phospho-STAT3 Tyr705 (Cell Signaling Technology, 9145S), MYL9 (Proteintech, 15354-1-AP), ACKR3/CXCR7 (Proteintech, 60216-1-Ig), CXCL8 (Proteintech, 31533-1-AP), CXCL12 (Proteintech, 17402-1-AP), CXCR4 (Proteintech, 60042-1-Ig), SMI32 (BioLegend, 801702), APP (Proteintech, 25524-1-AP), SOX2 (Cell Signaling Technology, 23064S), GFAP (Cell Signaling Technology, 3670T), CD44 (Cell Signaling Technology, 3570S), Nestin (10C2; Cell Signaling Technology, 33475), RRM2 (Proteintech, 11661-1-AP), PDGFRA (Cell Signaling Technology, 3174S), Ki-67 (Proteintech, 66555-6-Ig), OLIG2 (Abcam, ab109186), EGFR (Cell Signaling Technology, 4267S), CD3 (Proteintech, 60181-1-Ig), CD68 (Proteintech, 25747-1-AP), IBA1 (Proteintech, 66827-1-Ig), ACTA2/α-SMA (Proteintech, 14395-1-AP), CD31 (Novus Biologicals, NB600-562), MBP (Santa Cruz Biotechnology, sc-271524), CA9 (Proteintech, 11071-1-AP), gelsolin (BioLegend, 866501), and YKL-40/CHI3L1 (Cell Signaling Technology, 47066S). Anti-mouse Alexa Fluor Plus 555 (Thermo Fisher Scientific, A32727) and anti-rabbit Alexa Fluor 647 (Thermo Fisher Scientific, A32733) were used for multiplex detection, with DAPI nuclear counterstaining.

Conventional immunofluorescence used CD44 clone 156-3C11 (Cell Signaling Technology, 3570), IRF9 (Proteintech, 14167-1-AP), NGFR/p75NTR (Proteintech, 55014-1-AP) and phospho-MLC2 Ser19 where indicated. Immunoblotting used phospho-MLC2 Ser19 (Cell Signaling Technology, 3675S), MYL9 (Proteintech, 15354-1-AP) and GAPDH (Proteintech, 10494-1-AP).

### Orthotopic RNA profiling and survival

Processed single-cell RNA counts were aggregated within each sample and treatment condition to generate sample-level pseudobulk libraries. The post-TMZ versus post-DMSO comparison shown in Fig. 7 comprised four independent mice per condition, with one sample-level pseudobulk generated per mouse, representing 11,138 post-DMSO cells and 15,655 post-TMZ cells in total. Low-expression genes were removed using edgeR filterByExpr with min.count = 10, followed by trimmed mean of M values (TMM) normalization. Differential expression was assessed using robust edgeR quasi-likelihood generalized linear models with treatment condition as the explanatory variable. Fig. 7 presents the post-TMZ versus post-DMSO contrast. Genes were ranked by sign(log₂ fold change) × −log₁₀(P value). Targeted gene-set enrichment was performed using GSEApy v1.3.0 with 20,000 permutations and predefined program memberships, with Benjamini–Hochberg correction across the ten displayed programs.

Survival experiments compared vehicle, TMZ, fasudil and TMZ plus fasudil in GBM38, GBM43 and GBM123 orthotopic models. Treatment began 7 days after intracranial implantation and continued for 10 consecutive days. Fasudil (50 mg kg⁻¹) and TMZ (2.5 mg kg⁻¹) were administered intraperitoneally once daily, with combination-treated mice receiving both agents on the same schedule. Survival was summarized using Kaplan–Meier estimates and median survival. Treatment effects across models were evaluated using model-stratified Cox regression and stratified log-rank tests.

### Statistical reporting

GSEA q values, gene-level FDR values and adjusted P values are reported separately. Human between-group GSEA retained the corresponding prespecified full testing families. For the bulk RNA-seq analysis in Fig. 6, pathway q values retained Benjamini–Hochberg correction across the original 11-program family within each contrast, irrespective of the four programs selected for display. For the targeted orthotopic transcriptional analysis in Fig. 7, pathway FDR values were Benjamini–Hochberg adjusted across the ten displayed programs. Within-patient summaries and conditional spatial-bootstrap intervals were interpreted at their respective sampling levels. Nominal protein-program P values and image-field bootstrap intervals were not treated as independent-patient or independent-culture validation. Hazard ratios and 95% confidence intervals were estimated using model-stratified Cox regression, whereas P values for survival comparisons were obtained from stratified log-rank tests. Where field-level statistical comparisons are displayed, they quantify technical within-experiment consistency and were not interpreted as evidence of independent biological replication.

## Data and code availability

Data supporting the findings of this study are available from the corresponding author upon reasonable request. Processed datasets and analysis code underlying the principal analyses will be deposited in public repositories before journal publication.

## Funding

This work was supported by National Institute of Neurological Disorders and Stroke grants 1R01NS096376 and 1R01NS112856, National Cancer Institute grant P50CA221747 (SPORE for Translational Approaches to Brain Cancer), and a Brain Up Foundation grant (to A.U.A.).

## Competing interests

The authors declare no competing interests.

## Author contributions

R.C. and A.U.A. conceived and designed the study. R.C. performed computational analyses, integrated the datasets, prepared the figures and drafted the manuscript. U.H.F., P.U.K., N.B.D., L.K., J.J., J.R.T., M.C.O., V.F., A.A., S.W., J.C., S.A., H.K., B.A., P.J., K.M., A.M.C., M.C.T., R.S., M.S.L., J.M., K.S.D., D.P., D.H.H. and A.M.S. contributed to data generation, experimental work, clinical material, analysis, interpretation or study resources, as appropriate. A.U.A. supervised the study. R.C. and A.U.A. revised the manuscript with input from all authors. All authors reviewed and approved the final manuscript.

---

**Supplementary Figure 1.**
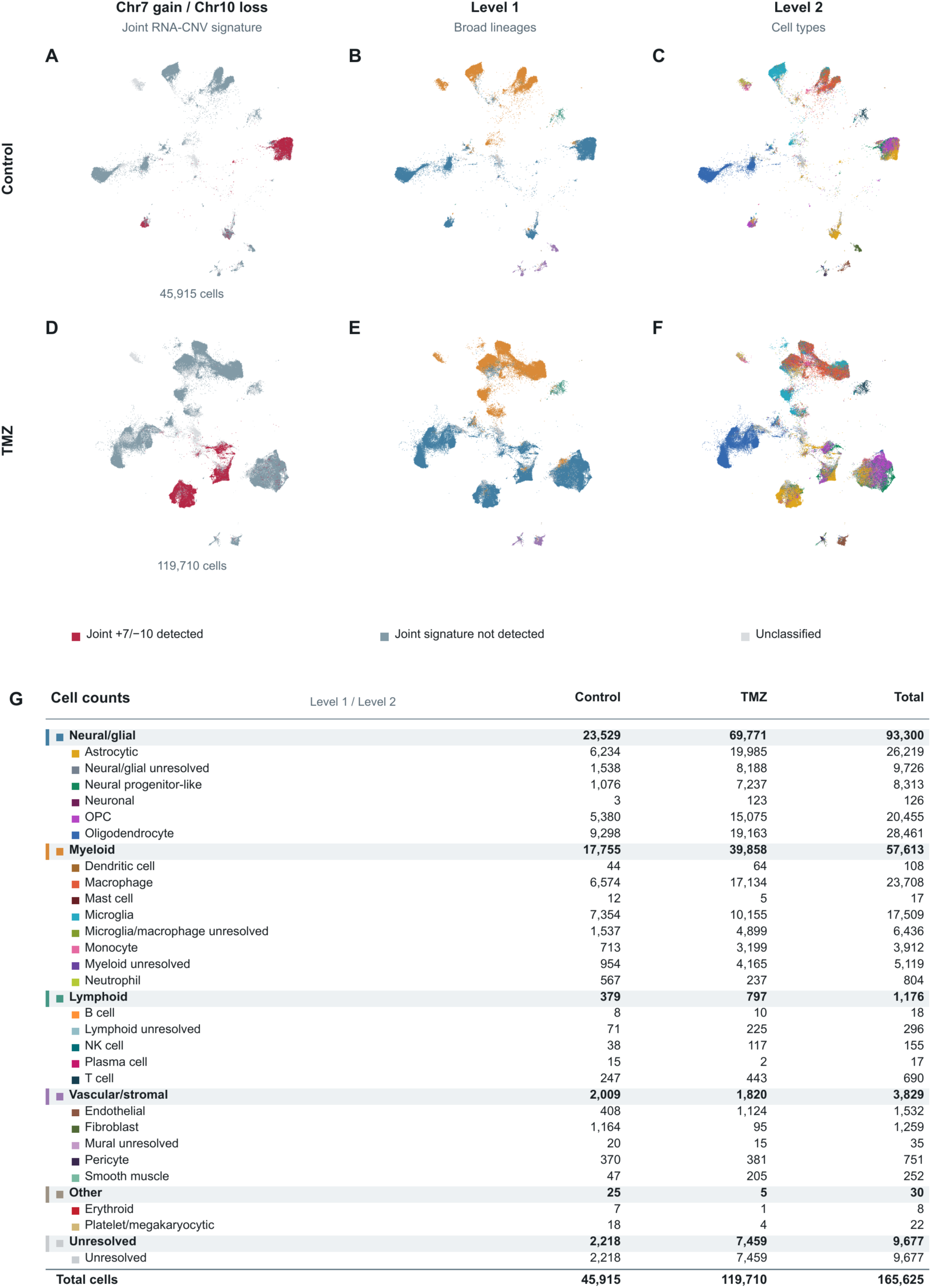
Hierarchical cellular annotation and RNA-inferred copy-number status in recurrent glioblastoma. **A**–**F**, Single-cell embeddings of recurrent comparators without additional preoperative temozolomide (TMZ; **A**–**C**, 45,915 cells) and tumors receiving additional preoperative TMZ (**D**–**F**, 119,710 cells). **A**,**D**, Joint RNA-inferred chromosome 7 gain/chromosome 10 loss signature. Red indicates detection of the joint signature, blue-gray indicates that the joint signature was not detected, and light gray denotes unclassified cells. **B**,**E**, Broad-lineage annotations (Level 1). **C**,**F**, Finer cell-type annotations (Level 2). **G**, Cell counts by broad lineage and cell-type annotation in each treatment group and across the combined dataset of 165,625 cells. Colors identify the corresponding annotation categories; unresolved assignments are retained explicitly.

**Supplementary Figure 2.**
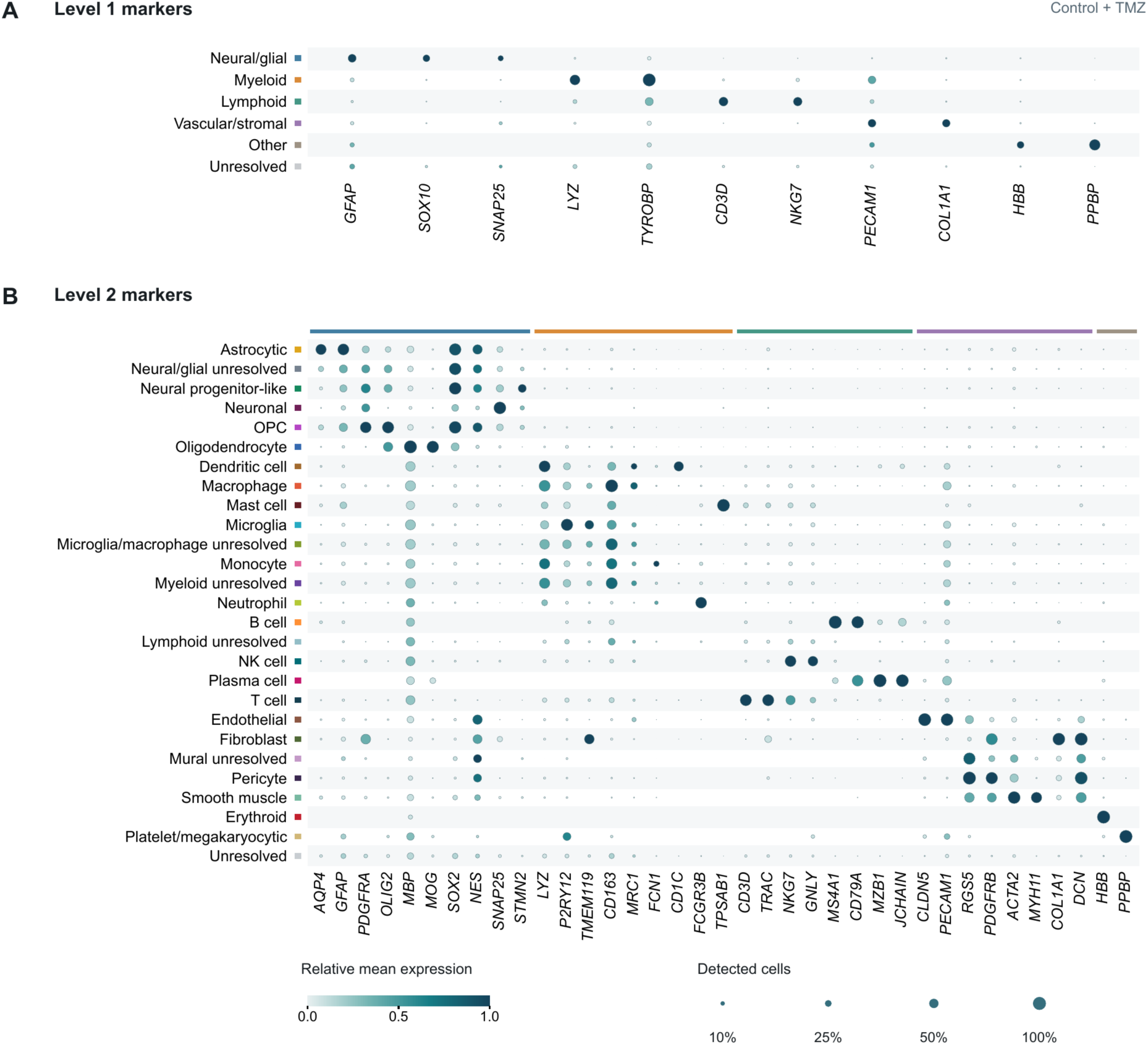
Marker-expression profiles supporting hierarchical single-cell annotation. **A**, Dot plot of selected marker genes across broad-lineage annotations (Level 1), including neural/glial, myeloid, lymphoid, vascular/stromal, other and unresolved populations. **B**, Dot plot of selected marker genes across finer cell-type annotations (Level 2), including unresolved subgroups. Rows represent annotated populations and columns represent genes. Dot size indicates the percentage of cells with detectable expression, and color indicates relative mean expression on the displayed 0–1 scale. Data combine the comparator and additional-preoperative-TMZ groups. Annotation colors correspond to those used in Supplementary Fig. 1. OPC, oligodendrocyte precursor cell; NK, natural killer.

**Supplementary Figure 3.**
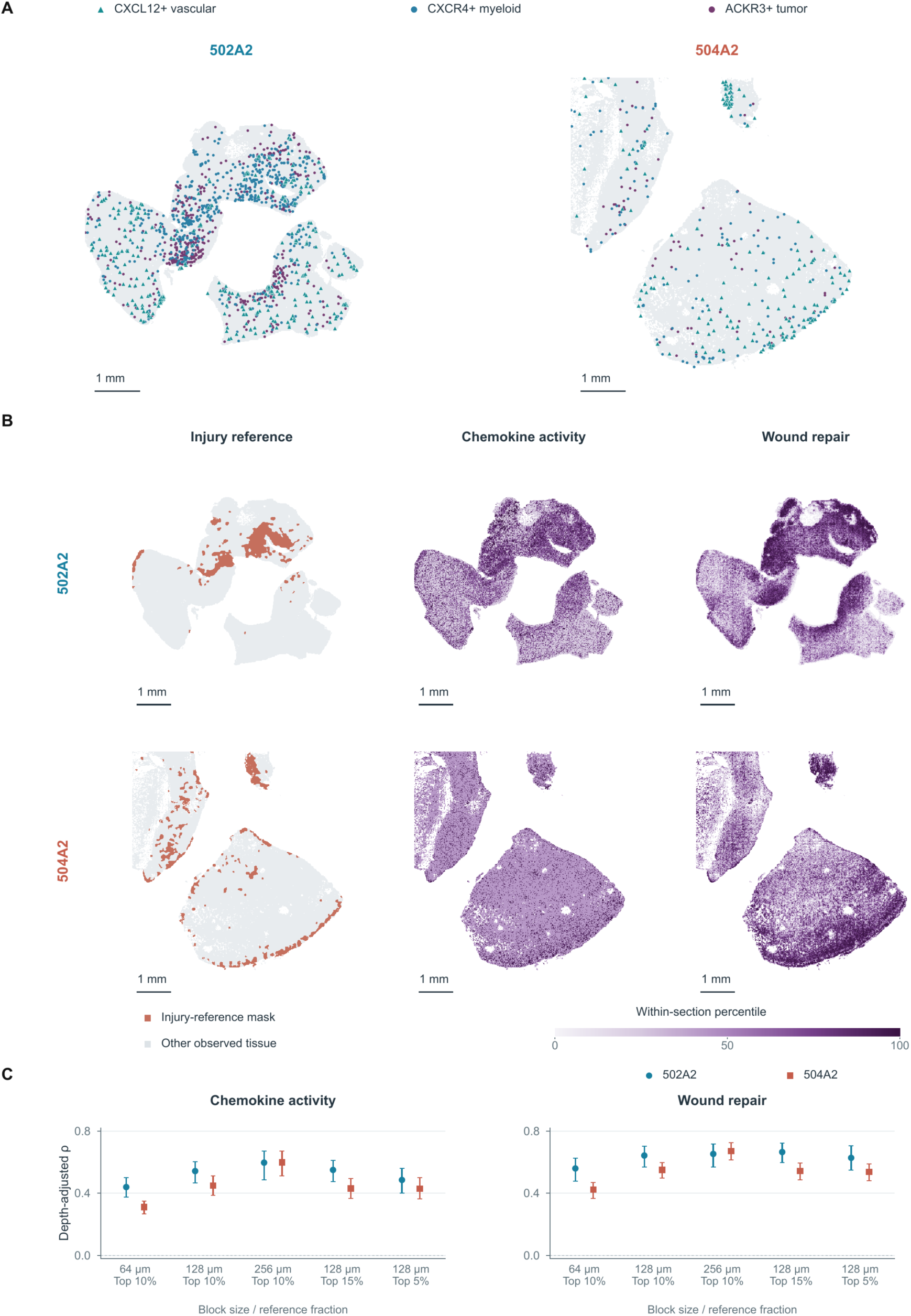
Native-resolution spatial context and robustness of injury-associated transcription. **A**, Native 16-µm maps of CXCL12-positive vascular, CXCR4-positive myeloid and ACKR3-positive tumor-enriched bins in TMZ-exposed sections 502A2 and 504A2; gray shows other retained tissue bins. **B**, RNA-defined injury-reference masks (top 10%, with connected-component filtering), chemokine activity and wound repair in the same sections. Program colors represent within-section percentiles, retaining tied values. Reference genes were excluded from the tested programs. Maps are unsmoothed, preserve tissue gaps and do not compare absolute expression between specimens. Scale bars in A,B, 1 mm. **C**, Whole-tissue injury-proximity correlations across 64-, 128- and 256-µm blocks using the top-10% reference, and 128-µm blocks using top-15% and top-5% references. Points show partial rank correlations adjusted for mean log library depth and mean log detected features; whiskers indicate conditional 95% spatial-bootstrap intervals from 500 resamples of 512-µm macroblocks, with nuisance fits held fixed. These models include reference interiors and differ from the composition- and tissue-edge-adjusted, outside-reference analyses in Fig. 2.

**Supplementary Figure 4.**
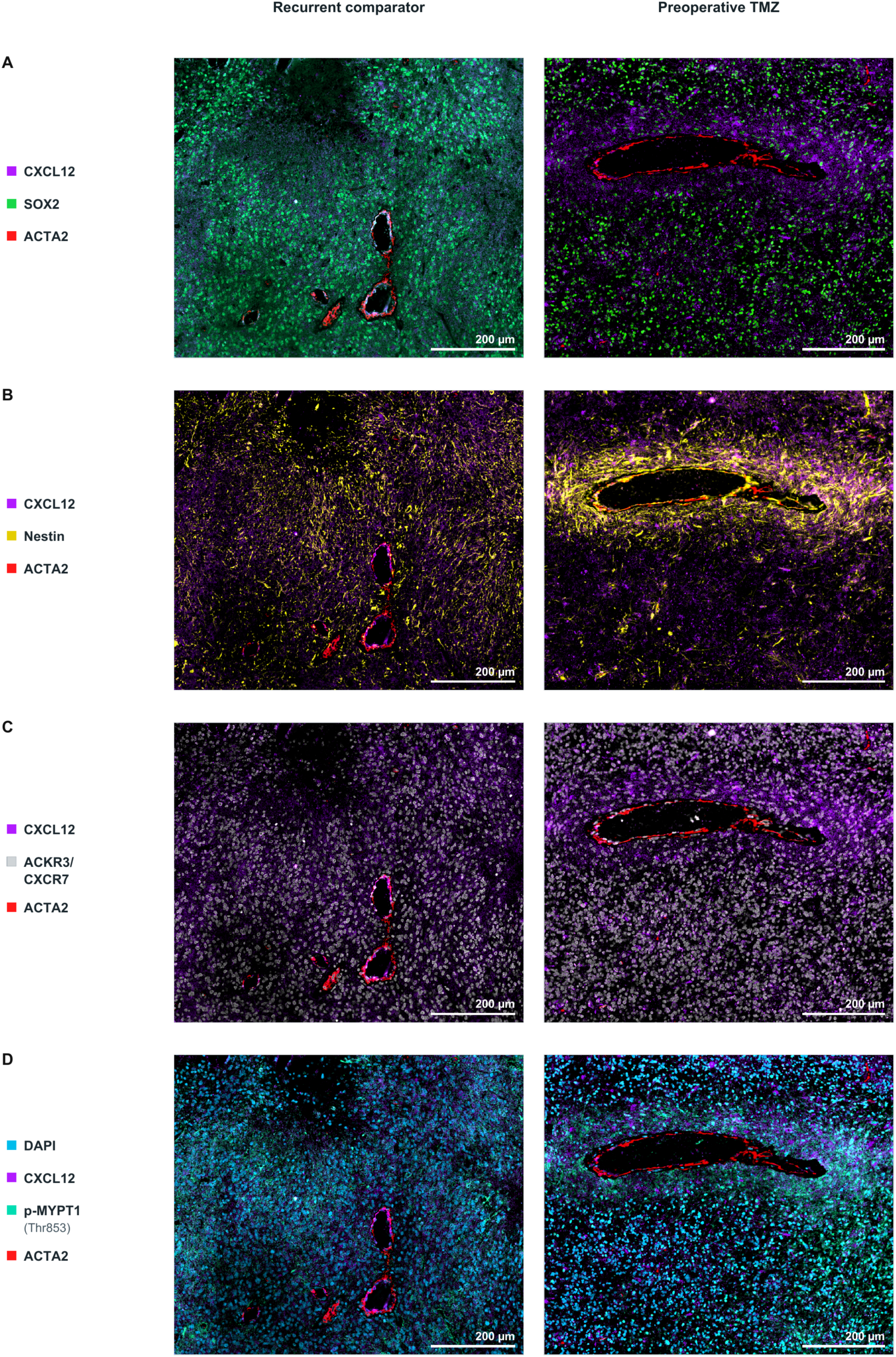
Multiplex tissue context for CXCL12 and glioblastoma-associated markers. **A**–**D**, Representative multiplex tissue images from a recurrent comparator without additional preoperative TMZ (left column) and a preoperative-TMZ specimen (right column). **A**, CXCL12, SOX2 and ACTA2. **B**, CXCL12, Nestin and ACTA2. **C**, CXCL12, ACKR3/CXCR7 and ACTA2. **D**, DAPI, CXCL12, p-MYPT1 (Thr853) and ACTA2. The same field is shown across rows within each specimen and corresponds to the field in Fig. 3E. CXCL12, purple; SOX2, green; Nestin, yellow; ACKR3/CXCR7, white; ACTA2, red; p-MYPT1 (Thr853), green/teal; DAPI, cyan-blue. Scale bars, 200 µm.

**Supplementary Figure 5.**
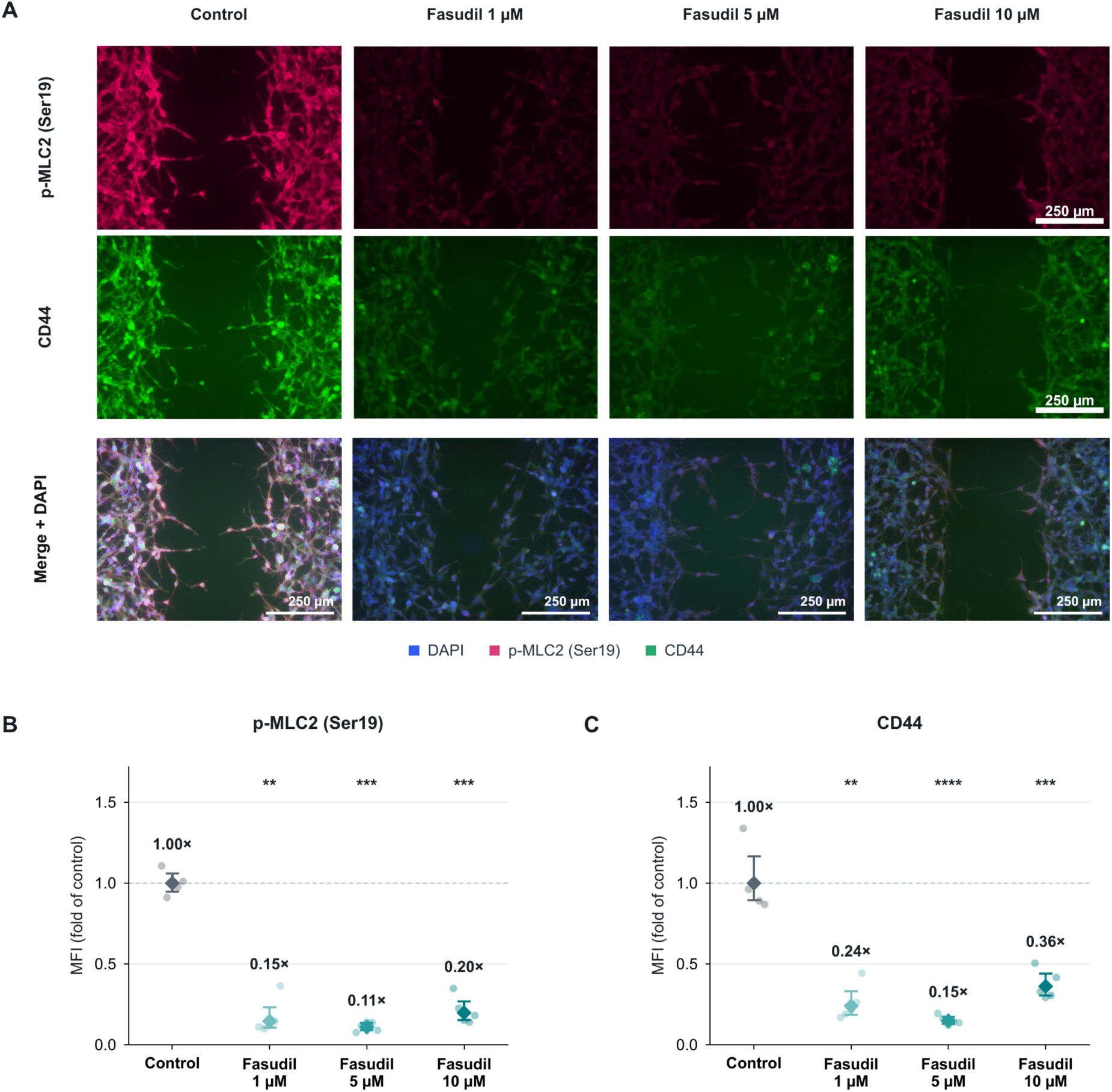
Fasudil-associated reductions in p-MLC2 and CD44 fluorescence. **A**, Representative immunofluorescence images of GBM43 cells under control conditions or following fasudil treatment at 1, 5 or 10 µM. p-MLC2 (Ser19), magenta; CD44, green. Marker-isolated views show the same representative field within each condition, using the source display settings. Scale bars, 250 µm. **B**,**C**, Background-subtracted mean fluorescence intensity of p-MLC2 (Ser19; B) and CD44 (C) in a DAPI-derived cellular proxy mask, normalized to the control geometric mean. Small circles represent five image fields from one stained well per condition. Diamonds and whiskers show the geometric mean and technical 95% bootstrap interval from 40,000 field resamples. The dashed line denotes control = 1. For the displayed technical comparisons, two-sided paired t tests on log₂-transformed field intensities were performed, with fields paired by position and Holm correction across the three control comparisons within each marker. **P < 0.01, ***P < 0.001 and ****P < 0.0001.

## Notes

### Competing Interest Statement

The authors have declared no competing interest.

